# Reimagining productive chemical space for RNA recognition beyond aromaticity

**DOI:** 10.64898/2026.08.10.743988

**Authors:** Lukasz T. Olenginski, Dia Patel, Aleksandra J. Wierzba, Robert T. Batey

## Abstract

Contemporary RNA-binding ligand collections are biased toward aromatic scaffolds, although it remains unclear whether this over-representation reflects an intrinsic requirement for productive RNA recognition or historical discovery bias. Here, using a modular “host-guest” ligand design strategy targeting the *env8* cobalamin (Cbl) riboswitch, we established a common molecular framework to directly evaluate whether aromaticity is fundamentally required for RNA binding. We synthesized a focused series of cyclic aliphatic β-axial Cbl derivatives, expanding the ligand library and enabling matched-pair comparisons to isolate the contribution of aromaticity to molecular recognition. Aliphatic ligands supported high-affinity RNA binding and regulatory activity comparable to aromatic analogues, with several derivatives exhibiting equal or greater affinity than their matched aromatic counterparts. Structural analyses revealed that aromatic and aliphatic ligands engage the same cryptic RNA binding site through distinct modes of molecular recognition, including nucleobase π-stacking and alternative van der Waals packing arrangements. Machine learning analyses further demonstrated that the physicochemical features associated with affinity extend beyond aromaticity itself and instead reflect a broader combination of shape, surface, heteroatom, and electronic properties. Together, these findings demonstrate that high-affinity RNA binding can arise from multiple structural and physicochemical solutions, suggesting that aromaticity is not uniquely privileged as a strategy for RNA-targeted ligand design and supporting broader exploration of underrepresented RNA-binding chemotypes.

## Introduction

The transcriptome has emerged as an increasingly important therapeutic target, driven by advances in RNA biology and a growing appreciation for the central roles of these biomolecules in regulating cellular function and disease.^1–3^ Recent successes in the discovery of RNA-targeting ligands have demonstrated that small molecules can selectively modulate transcript structure and function, establishing RNA as a viable target for chemical biology and drug discovery.^4–7^ Despite this progress, the principles governing productive molecular recognition remain comparatively underdeveloped relative to those established for proteins.^8,9^ Decades of medicinal chemistry and structural biology have produced extensive design frameworks for protein-targeted ligands, whereas the chemical space capable of engaging RNA remains poorly defined.^10^

To address this challenge, considerable effort has been devoted to identifying the physicochemical principles underlying RNA recognition. RNA-binding repositories (ROBIN,^11^ InfoRNA,^12^ R-BIND,^13^ and DRTL^14^) together with RNA-focused screening collections and comparative cheminformatic analyses, have provided important insights into the characteristics of RNA-binding chemical matter and established a foundation for RNA-targeted discovery.^11–17^ Nevertheless, many features repeatedly associated with productive binding have yet to be rigorously examined in a controlled experimental system, making it difficult to distinguish fundamental recognition principles from historical discovery bias.

Aromatic functionality is widely regarded as an important contributor to RNA recognition because planar aromatic systems can engage nucleobases through favorable π-stacking interactions.^18–20^ Consequently, aromatic scaffolds have become a prominent feature of many reported RNA-binding ligands and have strongly influenced contemporary RNA-targeted medicinal chemistry. However, because π-stacking exploits a general feature of RNA structure, such interactions may predispose aromatic-rich ligands toward broader transcriptome-wide recognition and present challenges for achieving target selectivity (**Figure 1A,B**).^4–7^ These considerations motivate the exploration of alternative chemotypes capable of engaging RNA through orthogonal recognition strategies. Among these, sp^3^-rich aliphatic scaffolds offer access to chemically distinct molecular topologies that may promote greater shape complementarity with local RNA pockets (**Figure 1C,D**) while simultaneously expanding the chemical space available for RNA-targeted ligand discovery. Whether such chemotypes can support productive recognition as effectively as aromatic ligands remains largely unexplored.

**Figure 1.**
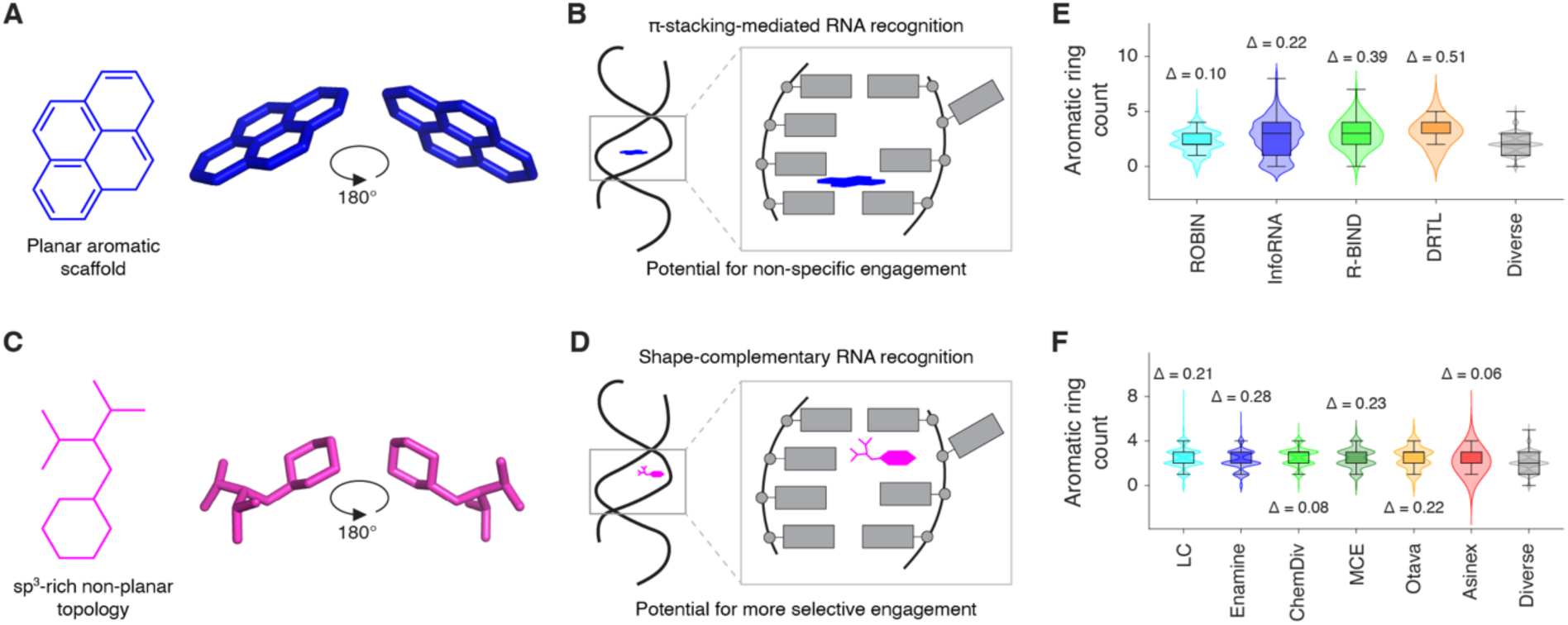
Conceptual framework and aromatic enrichment within contemporary RNA-binding chemical space. (A) Representative aromatic scaffold illustrating a topology commonly observed among RNA-binding ligands. (B) Conceptual representation of π- stacking-mediated RNA recognition facilitated by extended aromatic surfaces. (C) Representative aliphatic scaffold illustrating an alternative molecular topology. (D) Conceptual representation of an alternative recognition strategy in which non-planar chemotypes engage local RNA pockets through shape complementarity. Panels A-D are schematic illustrations and are not derived from experimental structural data. (E) Violin plots showing the distribution of aromatic ring counts among publicly available RNA-binding ligand repositories (ROBIN,^11^ InfoRNA,^12^ R-BIND,^13^ and DRTL^14^) relative to a chemically diverse reference collection from ChemDiv (“Diverse”). Values above each distribution indicate Cliff’s delta effect sizes (Δ) relative to the diverse set. (F) Distribution of aromatic ring counts among commercially available RNA-focused screening collections, including Life Chemicals (LC), Enamine, ChemDiv, MedChemExpress (MCE), Otava, and Asinex, in reference to the same diverse set.

Addressing this question experimentally is challenging. Reported RNA-binding ligands differ substantially in scaffold architecture, molecular size, charge, flexibility, and functional group composition,^11–14^ making it difficult to isolate the contribution of aromaticity to molecular recognition. Consequently, comparisons between aromatic and non-aromatic chemotypes are frequently confounded by broader differences in physicochemical properties. Resolving this question requires a system in which different chemotypes can be examined in a common molecular and structural context.

Previously, we developed a modular “host-guest” ligand design strategy targeting the *env8* cobalamin (Cbl) riboswitch in which chemically diverse “guests” are installed at the β-axial position of a conserved Cbl “host” (**Figure 2A,B**).^21,22^ In this framework, Cbl serves as a soluble host scaffold, while the β-axial substituent functions as a chemically tunable guest that is presented directly to the RNA binding pocket.^21^ This architecture enables systematic diversification of ligand chemistry while preserving a common molecular scaffold and recognition framework. We previously established that aromatic, aliphatic, and aromatic-aliphatic hybrid guest chemotypes can be accommodated within the *env8* binding site and that modification of the displayed guest fragment can modulate RNA affinity and function (**Figure 2B, S1**).^21,22^ The platform has additionally enabled high-resolution structural characterization of ligand recognition, which revealed that derivative binding occurs through β-axial engagement of a cryptic RNA binding site formed through ligand-induced base displacement (**Figure 2C, S2**).^21^

**Figure 2.**
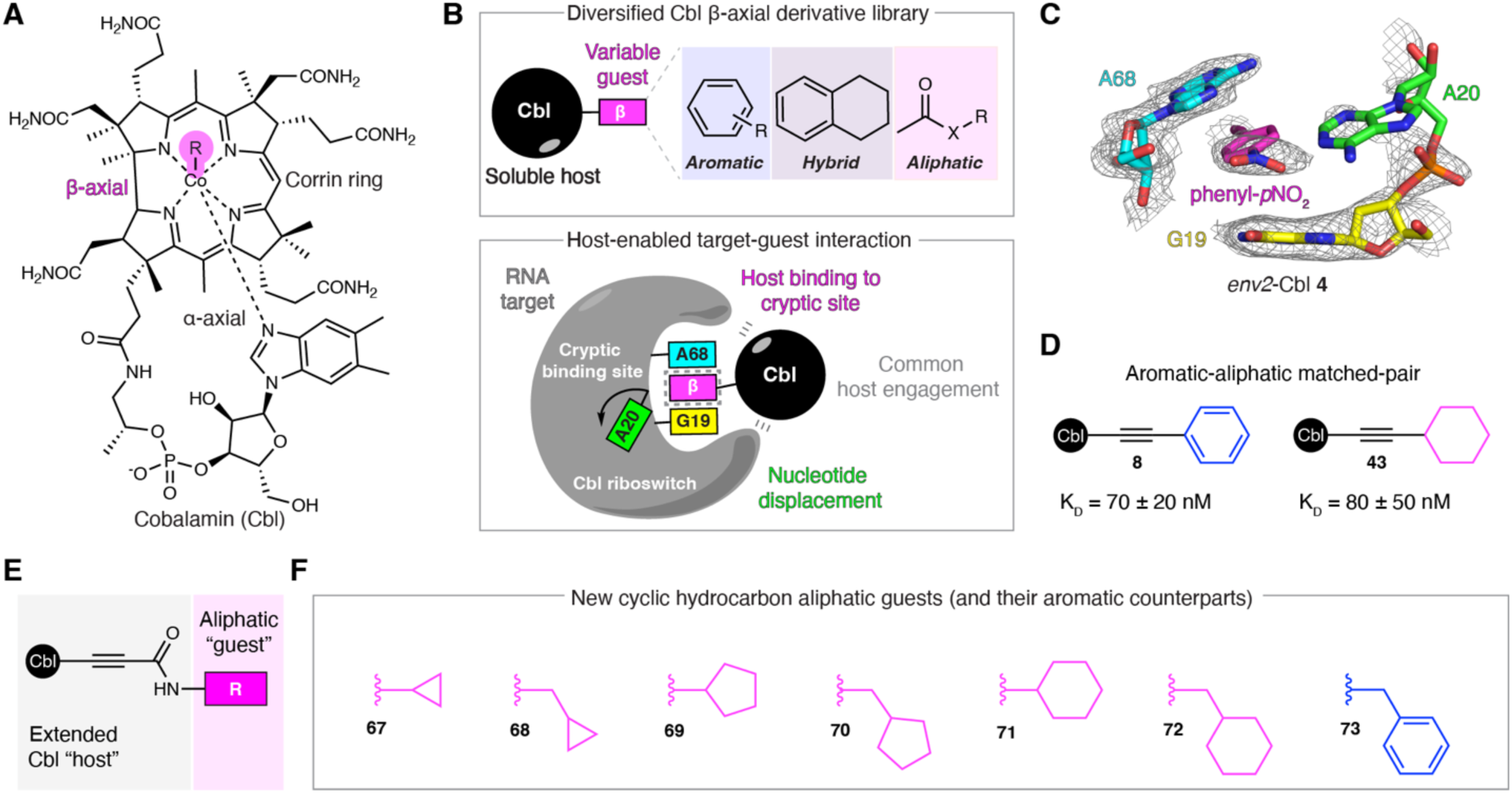
The *env8* cobalamin (Cbl) riboswitch model system for evaluating aromatic and aliphatic RNA recognition strategies. (A) Structure of Cbl highlighting the β-axial position used for ligand diversification. (B) Schematic overview of the “host-guest” platform.^21^ A common Cbl scaffold (“host”) displays variable β-axial substituents (“guests”) to a local RNA binding pocket, enabling systematic evaluation of how aromatic, aliphatic, and aromatic-aliphatic hybrid chemotypes influence RNA recognition within an otherwise conserved molecular framework. (C) Previously determined crystal structure of phenyl- *p*NO_2_-Cbl (**4**) (PDB 9E5I^21^) bound to the homologous *env2* Cbl riboswitch aptamer domain. Binding of Cbl **4** displaces A20 from the RNA core, exposing a cryptic binding site occupied by the β-axial group.^21^ In all structural representations, binding-pocket nucleotides G19 (yellow), A20 (green), and A68 (cyan) are numbered according to the full-length *env8* riboswitch^26^ and colored accordingly; β-axial substituents are shown in magenta. Mesh representations correspond to simulated-annealing 2F_o_-F_c_ omit maps in which A20 and the ligand were omitted from the model and are contoured at 1σ. (D) Previously reported aromatic-aliphatic matched-pair comparison demonstrating comparable RNA-binding affinities for phenyl- and cyclohexyl-containing derivatives.^21^ Experimentally determined K_D_ values are shown.^21^ (E) General structure of the extended amide-linked β-axial scaffold used to access additional aliphatic chemotypes. (F) Structures of the newly synthesized cyclic hydrocarbon derivatives (**67-72**) and aromatic comparator **73** evaluated in the present study.

Here, we compared contemporary RNA-binding chemical matter with representative chemically diverse small molecules and identified aromaticity as one of the most prominent distinguishing features of reported RNA-binding ligands. We then used the *env8* Cbl riboswitch to directly evaluate whether this enrichment reflects a fundamental requirement for productive RNA recognition by expanding the β-axial derivative library with a series of cyclic aliphatic fragments and leveraging matched-pair ligand comparisons. Through quantitative binding and functional measurements, structural characterization, and cheminformatic modeling, we demonstrate that aromatic chemotypes are not uniquely privileged for productive RNA recognition and that high-affinity binding can arise from multiple structural and physicochemical solutions. Collectively, these findings expand the chemical space capable of engaging RNA and provide a broader framework for RNA-targeted ligand design.

## Results and discussion

### Contemporary RNA-binding chemical space is biased toward aromatic scaffolds

To place our experimental studies in the context of contemporary RNA-targeted ligand discovery, we compared four literature-derived RNA-binding repositories (ROBIN,^11^ InfoRNA,^12^ R-BIND,^13^ and DRTL^14^) together with six commercially available RNA-focused screening collections against a chemically diverse reference library using a panel of seven physicochemical descriptors capturing major aspects of ligand composition and molecular architecture (**Figure S3, Tables S1-S6**). Across the RNA-binding repositories, aromatic ring content emerged as one of the most prominent distinguishing features relative to chemically diverse compounds, with Cliff’s Δ values ranging from 0.10-0.51 (**Figure 1E**). A similar, although less pronounced, trend was also observed among commercially available RNA-focused libraries (Δ = 0.06-0.28) (**Figure 1F**), indicating that aromatic enrichment extends beyond historical RNA-binding ligands to compound collections intentionally assembled for RNA-targeted discovery.

### Expansion of β-axial chemical space enables direct aromatic-aliphatic comparisons

The aromatic enrichment identified above (**Figure 1**) prompted us to systematically expand and re-examine our previously reported β-axial Cbl ligand collection, which comprises aromatic, aliphatic, and aromatic-aliphatic hybrid guest chemotypes (**Figure 2B, S1**).^21,22^ Within this dataset, the phenyl derivative **8** and the cyclohexyl derivative **43** exhibited nearly identical *env8*-binding affinities despite replacement of a planar aromatic group with its sp^3^-rich aliphatic counterpart (**Figure 2D**).^21^ Structural studies of phenyl-containing derivatives have established that aromatic guests engage the cryptic binding site through π-stacking interactions with A68 (**Figure 2C, S2**).^21^ In contrast, the cyclohexyl **43** lacks the planar aromatic surface required for π-stacking interactions and is therefore expected to engage A68 primarily through van der Waals contacts while likely occupying a similar region of the cryptic site. The nearly identical binding affinities of **8** and **43** therefore suggest that replacement of a nucleobase π-stacking interaction with aliphatic van der Waals contacts can support comparable energetics within this binding environment. Additional structures of aliphatic β-axial derivatives similarly demonstrate productive recognition through interaction modes distinct from nucleobase π-stacking within the same binding site (**Figure S2**).^21^ Together, these observations motivated a more systematic exploration of aliphatic chemical space.

Several considerations guided selection of the new β-axial ligands. First, multiple aliphatic derivatives within the original ligand set displayed relatively strong binding affinity and repressive activity,^21,22^ indicating that productive recognition of a pocket dominated by aromatic nucleobases could be achieved without extensive aromatic character. Second, amide-containing β-axial derivatives provided a synthetically accessible^23,24^ and structurally tractable^21^ framework for diversification (**Figure S4A**). In this design, the amide serves as an extension of the Cbl host while the terminal substituent functions as the displayed guest fragment presented to the *env8* cryptic pocket (**Figure 2E**). We therefore synthesized a series of new amide-linked derivatives incorporating cyclic hydrocarbon guests with and without a methylene spacer (**67**-**72**) together with a corresponding aromatic analogue (**73**) (**Figure 2F**). This strategy simultaneously expanded the representation of aliphatic chemotypes within the β-axial ligand library while generating new aromatic-aliphatic matched pairs for systematic comparison of RNA affinity, function, and molecular recognition.

### Aliphatic β-axial derivatives support high-affinity binding and functional RNA regulation

The new amide-linked β-axial derivatives were first evaluated using a previously established Cbl fluorescence displacement assay,^21,22^ enabling direct comparison with the original ligand collection^21,22^ and integration into the expanded dataset. All newly synthesized derivatives bound the *env8* riboswitch with nanomolar affinity, with the aliphatic ligands (**67**-**72**) exhibiting dissociation constant (K_D_) values ranging from ∼20-40 nM (**Figure 3A, Table S7**). Notably, each of the aliphatic derivatives bound more tightly than the aromatic analogue **73**, which displayed substantially weaker affinity (K_D_ = 110 ± 30 nM) (**Figure 3A**).

**Figure 3.**
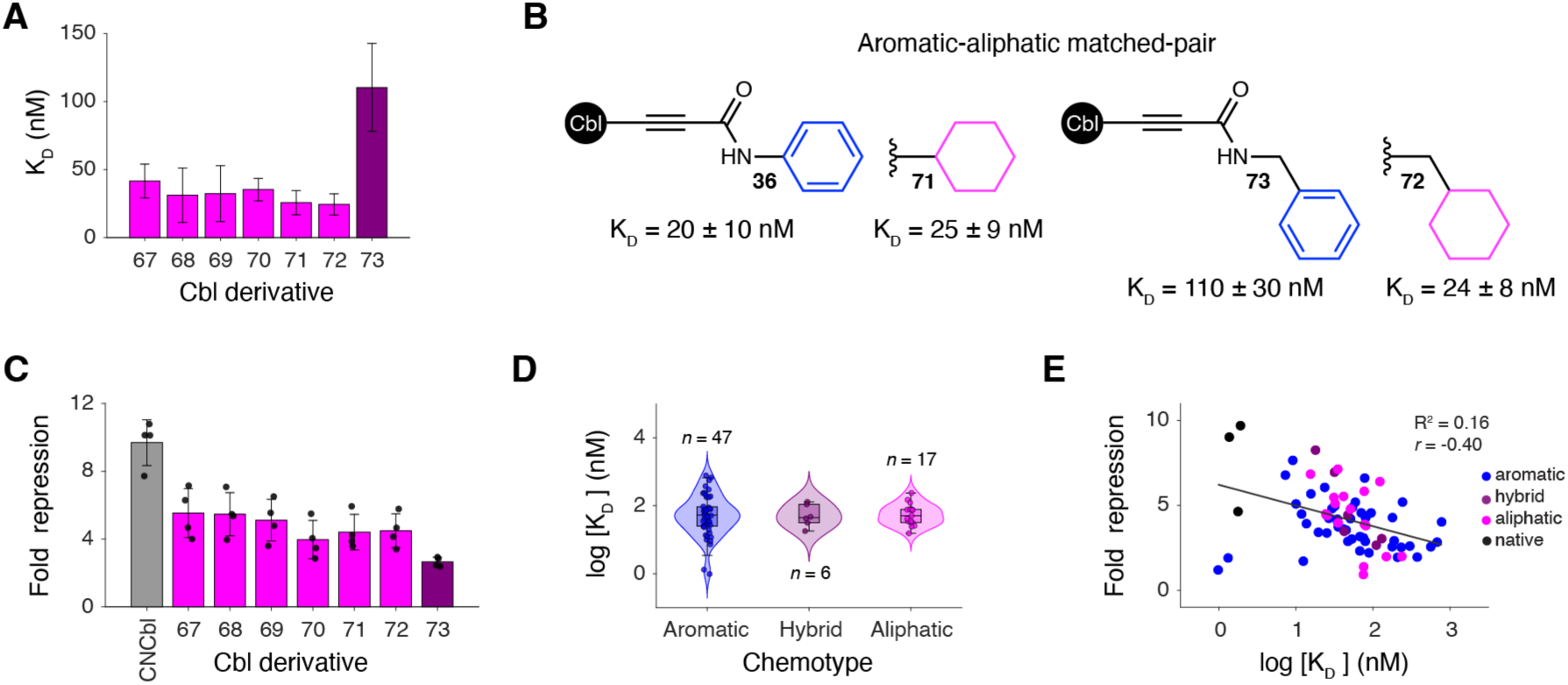
Aliphatic chemotypes support high-affinity RNA binding and riboswitch regulation. (A) K_D_ values for newly synthesized amide-linked aliphatic Cbl derivatives (**67- 72**) and aromatic comparator **73** determined by competitive fluorescence displacement assays. Data represent mean ± SD from three independent experiments (*n* = 3). (B) RNA- binding affinities of aromatic-aliphatic matched pairs used to directly assess the contribution of aromaticity to RNA recognition. (C) Functional activity of new amide-linked ligands **67-73** using a cell-based reporter assay in *E. coli*.^25^ Fold repression values represent mean ± SD from biological replicates (*n* = 4). (D) Violin plots showing the distribution of log-transformed *env8* K_D_ values for aromatic, hybrid, and aliphatic chemotypes across the complete β-axial ligand library. (E) Relationship between RNA- binding affinity (log-transformed K_D_ values) and riboswitch regulatory activity (fold repression) across the complete ligand set. Pearson correlation coefficient (*r*) is shown.

Examination of the amide-linked series further revealed distinct structure-activity relationships (SAR) (**Figure S4**). Within the direct attachment series (**67**, **69**, and **71**), increasing guest van der Waals volume was strongly associated with improved affinity (R^2^ = 0.99, *r* = -0.99), consistent with improved guest engagement of the cryptic pocket (**Figure S4B**). Although this relationship was monotonic across the cyclic hydrocarbon series, the absolute energetic gains remained relatively modest, indicating that even relatively small aliphatic substituents support productive recognition of the cryptic binding site. In contrast, this relationship largely disappeared upon introduction of a methylene spacer between the amide and displayed guest (**68**, **70**, and **72**; R^2^ = 0.20, *r* = -0.45), suggesting that increased conformational flexibility alters how guest size contributes to molecular recognition (**Figure S4B**).

Comparison of matched hydrocarbon pairs further supported this interpretation. While introduction of the spacer improved affinity for the smallest cyclopropyl derivative, little benefit was observed for the larger cyclopentyl and cyclohexyl analogues (**Figure 3A**), suggesting that additional conformational flexibility is only advantageous when sufficient unoccupied pocket volume remains available for productive exploration. Energetic analysis relative to the unsubstituted amide reference (Cbl **57**) further revealed that all amide-displayed guests contributed favorably to RNA recognition, albeit to varying degrees (**Figure S4C**). Notably, despite clear differences among individual guest chemotypes, the overall energetic landscape remained relatively compressed, with even simple aliphatic substituents providing substantial gains in binding free energy relative to the unsubstituted amide. Together, these results demonstrate that multiple cyclic aliphatic chemotypes support high-affinity RNA binding.

Direct aromatic-aliphatic matched-pair comparisons provided a more stringent evaluation of how guest chemotype influences RNA recognition. Replacement of the aromatic phenyl group in **36** (20 ± 10 nM^21^) from the original amide-linked series with the new aliphatic cyclohexyl group in **71** (25 ± 9 nM) resulted in nearly identical binding affinities (**Figure 3B**). A second matched-pair generated through the expanded ligand set produced an even more striking result. Whereas the aromatic derivative **73** bound with a K_D_ of 110 ± 30 nM, its aliphatic cyclohexyl counterpart **72** bound approximately five-fold more tightly (24 ± 8 nM) (**Figure 3B**). Together, these comparisons demonstrate that aromatic functionality is not uniquely privileged for high-affinity recognition of the *env8* cryptic site and, in some contexts, can be replaced by aliphatic chemotypes with little energetic consequence or even improved binding.

The newly synthesized derivatives also retained measurable regulatory activity in a cell-based reporter assay,^25^ producing approximately four-to-six-fold repression (**Figure 3C, Table S8**). The addition of the new amide-linked derivatives enabled reassessment of the complete β-axial ligand library and the extent to which affinity varies as a function of ligand chemistry. When grouped according to chemotype, aromatic, aliphatic, and hybrid derivatives exhibited broadly overlapping affinity distributions (**Figure 3D**). Although several of the tightest binders were aromatic,^21^ aliphatic ligands occupied a similar affinity range, and no statistically significant differences were observed between chemotype classes. Comparison of RNA affinity and regulatory activity across the complete dataset further revealed only a weak relationship between these parameters (**Figure 3E**). While tighter binding generally trended toward increased repression, affinity explained only a small fraction of the observed functional variance (R^2^ = 0.16; *r* = -0.40), and several of the highest-affinity ligands exhibited weak regulatory activity.^21^ Together, these observations suggest that aromatic functionality is not broadly privileged for productive RNA recognition within this system. Moreover, differences in cellular activity were not strictly proportional to RNA-binding affinity, indicating that permeability and other physicochemical or cellular properties may also influence functional outcomes.

### Aliphatic ligands engage the cryptic pocket through adaptable recognition modes

To investigate how aliphatic chemotypes engage the cryptic RNA binding site, crystal structures were determined for the complete series of newly synthesized amide-linked derivatives (**67**-**72**) bound to the homologous *env2* Cbl riboswitch aptamer domain (**Figure 4, Table S9, S10**). In addition to enabling structural interpretation of the matched-pair analyses described above, the resulting dataset provides atomic-level insight into the interaction strategies used by aliphatic chemotypes to achieve productive RNA recognition.

**Figure 4.**
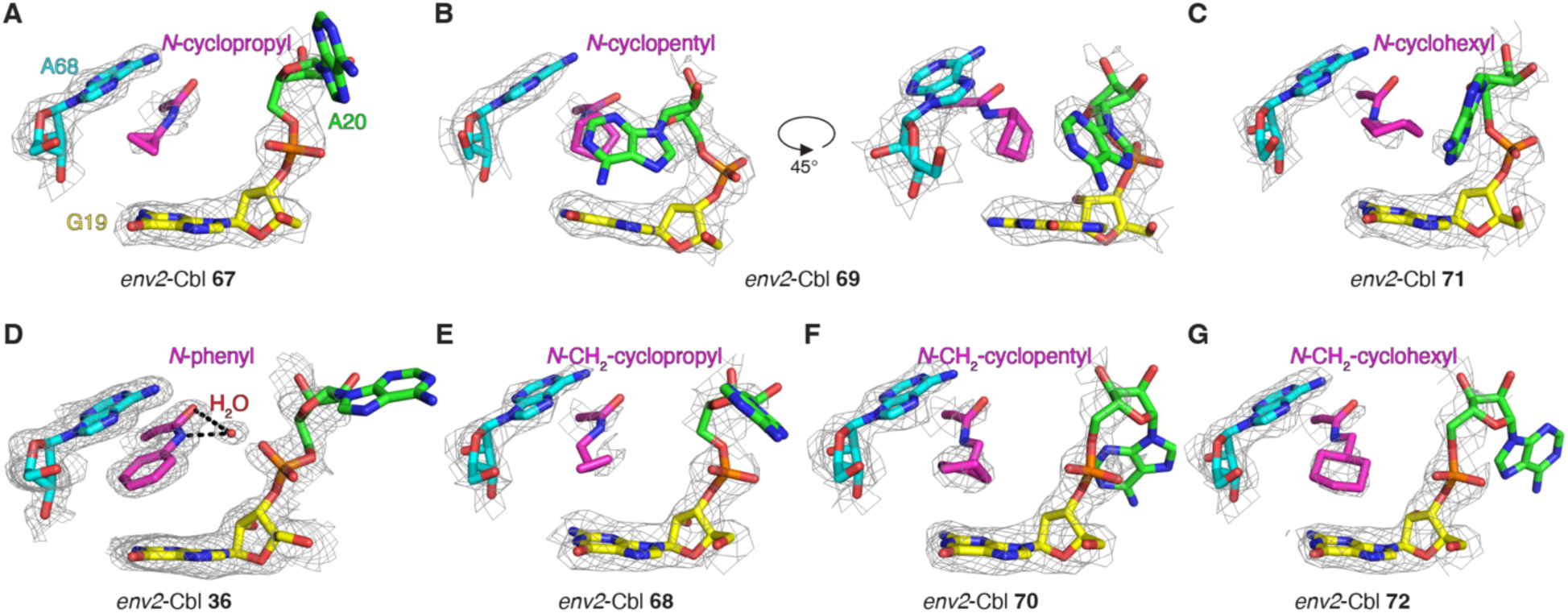
Structural basis for recognition of cyclic hydrocarbon aliphatic ligands within the cryptic binding site. Crystal structures of *N*-cyclopropyl (**67**) (A), *N*-cyclopentyl (**69**) shown in two views (B), *N-*cyclohexyl (**71**) (C), aromatic comparator *N*-phenyl (**36,** PDB 9E5Q^21^) (D), *N*-CH_2_-cyclopropyl (**68**) (E), *N*-CH_2_-cyclopentyl (**70**) (F), and *N*-CH_2_-cyclohexyl (**72**) (G) bound to the homologous *env2* Cbl riboswitch aptamer domain. Binding pocket nucleotides G19 (yellow), A20 (green), and A68 (cyan) are numbered according to the full-length *env8* riboswitch and colored accordingly; β-axial substituents are shown in magenta. Mesh representations correspond to simulated-annealing 2F_o_-F_c_ omit maps in which A20 and the ligand were omitted from the model and are contoured at 1σ. Dashed lines in panel D denote a water-mediated hydrogen-bonding observed for derivative **36**.^21^

Structural comparison of the phenyl- and cyclohexyl-containing derivatives reveals how distinct guest chemistries achieve comparable binding energetics (**Figure 4A-D**). Consistent with previous structures of phenyl-containing β-axial derivatives,^21^ the phenyl group of **36** is positioned beneath A68 where it participates in favorable π-stacking interactions with the nucleobase (**Figure 4D**). In contrast, the cyclohexyl-containing **71** adopts a distinct recognition mode in which the aliphatic ring is shifted toward G19 and engages the cryptic pocket primarily through van der Waals contacts (**Figure 4C**). Thus, despite exhibiting nearly identical RNA-binding affinities (K_D_ = 20 ± 10 nM^21^ and 25 ± 9 nM, respectively) (**Figure 3B**), the two ligands achieve recognition through different local interaction networks and distinct spatial solutions within the same binding site. Together with the energetic analysis (**Figure S4C**), these observations indicate that productive engagement of the cryptic pocket is not uniquely dependent on aromatic π-stacking interactions. Instead, both aromatic and aliphatic guest chemistries contribute favorably to RNA recognition relative to the unsubstituted amide reference (**57**), despite engaging the cryptic pocket through distinct local interaction networks.

Although structural interpretation of the second matched pair was limited by the absence of a co-crystal structure with **73**, the remaining aliphatic structures provided insight into the SAR trends observed across the amide-linked series (**Figure S4B**). The cyclopropyl derivatives occupied only a limited portion of the pocket, and introduction of a methylene spacer enabled the guest to extend further toward G19 while appearing to maintain interactions with A68 (**Figure 4A,E**). This observation provides a plausible structural rationale for the enhanced affinity of the spacer-containing cyclopropyl derivative (**Figure 3A**). In contrast, the larger cyclopentyl (**Figure 4B,F**) and cyclohexyl (**Figure 4C,G**) guests primarily extended toward G19 regardless of spacer incorporation, consistent with the comparatively modest energetic effects observed upon introduction of the methylene linker (**Figure 3A, S4B**). Together, these structures suggest that the energetic benefit of additional conformational flexibility depends on how effectively the displayed guest explores the available binding volume. Although the larger aliphatic guests occupy a greater portion of the cryptic pocket, they do not achieve optimal packing interactions, consistent with the modest energetic differences observed across the series. Beyond these general trends, the aliphatic structural series revealed recognition modes not observed among the aromatic derivatives. Most notably, the cyclopentyl derivative **69** adopted a markedly puckered conformation that enabled simultaneous engagement of G19 and a partially engaged A20 (**Figure 4B**). In contrast to the more fully displaced A20 observed in the other structures, this interaction mode preserved contacts with both nucleobases while maintaining productive occupation of the cryptic pocket. These observations illustrate how conformationally adaptable aliphatic guests can redistribute van der Waals contacts throughout the binding site and access alternative local interaction geometries. Collectively, these structural data demonstrate that productive engagement of the cryptic site can be achieved through multiple distinct recognition strategies. The ability of aliphatic guest chemistry to support high-affinity binding is particularly notable because formation of the cryptic pocket requires displacement of the A20 nucleobase, which normally occupies the site through aromatic π-stacking interactions. Rather than recapitulating this binding mode, the aliphatic ligands exploit distinct packing arrangements and conformational adaptability to achieve comparable energetic outcomes.

### Productive RNA recognition is associated with a multidimensional physicochemical framework

Having established that aromatic, aliphatic, and hybrid ligands all support productive recognition of the *env8* cryptic binding site, we next sought to identify the physicochemical features associated with RNA-binding affinity across the complete β-axial ligand library. Machine learning (ML) models were therefore constructed using experimentally measured binding affinities and molecular descriptors calculated for the β-axial groups of all ligands (**Figure 5A, S5A**). Random forest (RF) (**Figure 5D**) and gradient boosting (GB) (**Figure S5E**) models achieved improved predictive performance relative to elastic net (EN) regression (**Figure 5B**), suggesting that affinity is influenced by partially non-linear relationships among multiple molecular properties. However, because the primary objective was identification of reproducible physicochemical features associated with RNA binding rather than development of a predictive framework, subsequent analyses focused on EN models, which provide direct descriptor interpretability. Importantly, the descriptors identified by EN (**Figure 5C**) exhibited strong agreement with those recovered from RF feature-importance analyses (**Figure S5C,D, Tables S11**), indicating convergence on a common set of molecular features despite differences in model architecture.

**Figure 5.**
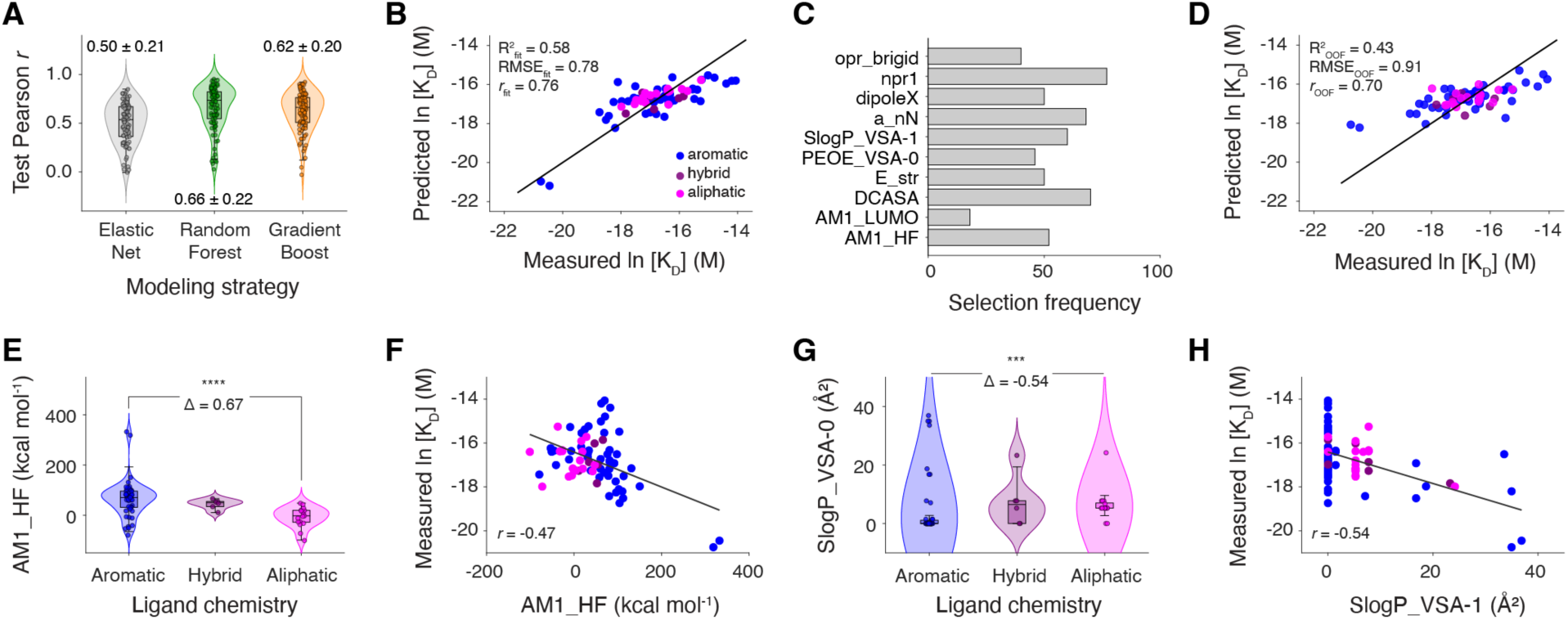
Machine learning (ML) identifies physicochemical features associated with *env8* binding beyond aromaticity alone. (A) Distributions of Pearson correlation coefficients (*r*) obtained from 100 repeated train-test iterations using elastic net (EN), random forest (RF), and gradient boosting (GB) regression models. Values above each distribution indicate mean ± SD. (B) Predicted versus experimentally measured binding affinity (natural log-transformed K_D_ values) for the final consensus EN model. Model performance statistics are shown. (C) EN descriptor selection frequencies across 100 modeling iterations. Descriptors selected in at least 30% of iterations were retained for construction of the final consensus model. (D) Out-of-fold (OOF) predictions from the RF model. Model performance statistics are shown. (E) Distribution of AM1_HF values across aromatic, hybrid, and aliphatic ligand classes. (F) Relationship between measured binding affinity ln(K_D_) and AM1_HF. (G) Distribution of SlogP_VSA-0 values across aromatic, hybrid, and aliphatic ligand classes. (H) Relationship between measured binding affinity ln(K_D_) and SlogP_VSA-0. Statistical significance was assessed using Wilcoxon rank-sum tests (***, p < 0.001; ****, p < 0.0001) and Cliff’s Δ effect size.

Across repeated cross-validation runs, EN consistently identified a compact set of descriptors associated with RNA-binding affinity (**Figure S5B**), including npr1, DCASA, a_nN, SlogP_VSA1, AM1_HF, and dipoleX (**Figure 5C, Table S12-S14**). These descriptors span molecular shape, exposed surface character, heteroatom content, hydrophobic surface partitioning, and electronic properties, suggesting that productive RNA recognition is governed by a multidimensional combination of physicochemical features rather than a single dominant molecular characteristic. Notably, all consensus descriptors exhibited complete sign consistency across model iterations, supporting a robust and reproducible relationship between these features and RNA engagement.

To determine how these features relate to β-axial guest chemistry, we compared descriptor distributions across aromatic, aliphatic, and hybrid chemotypes. Several descriptors strongly differentiated ligand class. Aromatic derivatives were enriched in descriptors associated with rigidity and π-electronic character, including opr_brigid (Cliff’s Δ = 0.85), AM1_HF (Δ = 0.67), and E_str (Δ = 0.62), whereas aliphatic ligands occupied distinct regions of descriptor space characterized by increased heteroatom content (a_nN, Δ = -0.41), altered surface-partitioning properties (SlogP_VSA1, Δ = -0.54), and distinct electronic profiles (AM1_LUMO, Δ = -0.83) (**Figures 5E,G, S6A, Table S12**). However, the descriptors most strongly associated with binding affinity only partially overlapped with those that most strongly differentiated chemotype classes. For example, a_nN, DCASA, SlogP_VSA1, npr1, and AM1_HF all exhibited measurable correlations with affinity (|*r*| = 0.47-0.59) (**Figure 5F,H, S6B**), yet these features were distributed across multiple descriptor classes and were not synonymous with aromaticity itself.

Collectively, these analyses indicate that productive recognition of the *env8* cryptic pocket is governed by a broader physicochemical framework than aromaticity alone. While aromatic and aliphatic ligands occupy distinct regions of chemical space, both chemotypes achieved high-affinity recognition. Together with the structural analyses, these findings suggest that aromatic and aliphatic ligands satisfy the requirements for productive RNA recognition through distinct but ultimately convergent combinations of molecular features. More broadly, these results support the view that high-affinity RNA binding can emerge from multiple regions of chemical space provided that the underlying physicochemical requirements for recognition are satisfied.

### Summary and Outlook

The historical dominance of aromatic scaffolds among reported RNA-binding ligands^11–14^ has fostered the widespread view that aromaticity is a principal determinant of productive RNA recognition. Using the *env8* Cbl riboswitch as a model system, we demonstrated that this relationship can be more nuanced. Through a combination of biophysical measurements, structural biology, and ML modeling, we show that aromatic functionality is not uniquely required for productive recognition of the *env8* cryptic binding site. In multiple matched-pair comparisons, direct substitution of aromatic functionality with aliphatic groups resulted in equivalent or improved RNA-binding affinity, demonstrating that productive recognition can be maintained across fundamentally different chemotypes.

A unifying observation emerging from these studies is that productive RNA recognition can be achieved through multiple solutions. Structurally, aromatic and aliphatic ligands engage the *env8* cryptic binding site through distinct local interaction networks, with the aliphatic ligands exploiting alternative packing arrangements and conformational adaptability to achieve comparable energetic outcomes. Physicochemically, the features associated with RNA-binding affinity extend beyond aromaticity itself and instead reflect a broader combination of molecular shape, surface character, heteroatom content, hydrophobic partitioning, and electronic properties. Together, these findings suggest that aromaticity should not be viewed as uniquely privileged among the strategies available for productive RNA recognition, but rather as one of several possible solutions by which ligands can satisfy the underlying requirements for binding. Expanding the repertoire of available molecular recognition strategies may ultimately provide additional opportunities to achieve selective RNA binding while broadening the medicinal chemistry toolbox available for RNA-targeted ligand discovery. More broadly, these results raise the possibility that contemporary RNA-targeted chemical space may be narrower than necessary. The historical emphasis on aromatic scaffolds has undoubtedly yielded many successful RNA-binding ligands.^11–14^ However, it may also have biased discovery efforts toward a limited subset of viable molecular solutions. Future studies will be required to determine the generality of these observations across additional RNA targets, ligand classes, and binding pockets. Nevertheless, the present findings suggest that broader exploration of underrepresented chemotypes may uncover productive recognition strategies that are not readily apparent from existing RNA-binding chemical matter, thereby providing new opportunities for hit discovery and subsequent ligand optimization.

## Materials and Methods

### RNA chemical space comparison

#### Analysis of Aromatic Ring Enrichment in RNA-Targeted Chemical Space

Publicly available RNA-binding ligand repositories (ROBIN,^11^ InfoRNA,^12^ R-BIND,^13^ and DRTL^14^), commercially available RNA-focused screening collections (Life Chemicals, Enamine, ChemDiv RNA, MedChemExpress, Otava, and Asinex), and a chemically diverse reference collection from ChemDiv were analyzed using Molecular Operating Environment (MOE) (Chemical computing group, v2022). Ligands were imported from SMILES strings (RNA-binding repositories) or supplier-provided SDF files (commercial libraries), curated to remove salts and/or counterions, assigned dominant protonation states at pH 7.4, and energy minimized in MOE using default parameters. Molecular descriptors were subsequently calculated from the minimized structures.

To provide a concise representation of molecular properties relevant to RNA recognition, seven descriptors spanning major physicochemical dimensions were selected for analysis: aromatic ring count (aromaticity; arorings), number of rotatable bonds (conformational flexibility; b_rotN), fraction rotatable bonds (relative flexibility; b_rotR), chiral centers (three-dimensionality; chiral), lipophilicity (logP (o/w))), relative polar surface area (polarity; relPSA), and molecular weight (molecular size; Weight). Aromaticity was quantified using the descriptor *arorings*, defined as the number of aromatic rings present within each molecule. Distributions of aromatic ring counts were visualized using violin plots and compared to the chemically diverse collection from ChemDiv (“Diverse”) using Cliff’s delta (Δ) effect sizes. Statistical significance was assessed using two-sided Wilcoxon rank-sum tests. Empirical cumulative distribution function (ECDF) analyses were performed to evaluate distributional shifts in aromatic ring content across datasets.

#### Synthesis of Cbl β-Axial Derivatives

Cbl β-axial derivatives **67**-**73** were synthesized using established synthetic procedures^23,24^ developed for β-axial Cbl functionalization. Full synthetic methods and are provided in the **Supporting Information**.

#### RNA Preparation

RNAs (sequences provided in **Table S15**) were prepared by *in vitro* transcription using PCR-amplified DNA templates and T7 RNA polymerase.^27^ Transcription products were purified by preparative denaturing polyacrylamide gel electrophoresis, buffer exchanged into Milli-Q H2O, and concentrated using centrifugal concentrators (Millipore-Sigma). RNA concentrations were determined by absorbance at 260 nm using extinction coefficients determined from the summation of the individual bases. Prior to binding experiments, RNAs were thermally annealed by heating at 95 °C for 2 min, cooling on ice for 10 min, and equilibrating to room temperature.

#### Competitive Fluorescence Displacement Assays

Binding affinities of Cbl derivatives were determined using a previously described competitive fluorescence displacement assay^21,22^ based on CNCbl-5xPEG-ATTO590.^28^ Briefly, 60 μL reactions contained 0.1 μM full-length *env8* RNA, 1 μM CNCbl-5xPEG- ATTO590, 1X RNA binding buffer (50 mM HEPES, pH 8.0, 100 mM KCl, 10 mM NaCl, and 1 mM MgCl_2_), and 0.01% Nonidet P40. Increasing concentrations of competing ligand were added and reactions were incubated for 1 h at room temperature. Following equilibration, samples were transferred to 384-well plates (Corning) and fluorescence was measured using a CLARIOstar Plus microplate reader (BMG Labtech) with excitation at 594 nm and emission collected from 620-670 nm. Fluorescence values were integrated across the emission spectrum and normalized to reactions lacking competitor. Displacement curves were fit in MATLAB (v2024b) using a single-site binding model with the Hill slope fixed to 1.0. IC_50_ values obtained from displacement experiments were converted to equilibrium dissociation constants (K_D_) using the previously determined affinity of CNCbl-5xPEG-ATTO590 for *env8*.^21,29^

#### Cell-Based Riboswitch Reporter Assays

Riboswitch regulatory activity was evaluated using a previously described *env8*-GFPuv reporter assay.^25^ Plasmids harboring *env8-*GFPuv (Addgene #99831^25^) or a pBR322 empty-vector control were transformed into *E. coli* JW3805 cells. For each biological replicate, 1 μL of a saturated overnight culture was inoculated into 1 mL of CSB medium supplemented with 100 μg/mL carbenicillin and 10 nM Cbl derivative and grown to mid-log phase at 37 °C. Aliquots (200 μL) from each biological replicate were transferred in technical triplicate to 96-well plates (Corning) and GFP fluorescence was measured using a Tecan Infinite 200 Pro microplate reader (395 nm excitation, 510 nm emission). Fluorescence values were normalized to cell density (OD_600_) and background-corrected using cells containing the empty-vector control. Fold repression values were calculated as the ratio of median normalized, background-corrected fluorescence measured in the absence and presence of ligand.^25^

#### Crystallization of *env2*-Ligand Complexes

The *env2* aptamer domain was crystallized in complex with Cbl derivatives using hanging-drop vapor diffusion at 30 °C. RNA-ligand complexes were prepared by combining *env2* RNA (250 μM) with Cbl derivative (375-500 μM) in 1X TE buffer. Crystallization drops containing 2 μL of RNA-ligand solution and 2 μL of precipitant solution (40 mM sodium cacodylate, pH 7.0, 10% (w/v) 2-methyl-2,4-pentanediol (MPD), 12 mM spermine tetrahydrochloride, 80 mM KCl, and 20 mM MgCl_2_) were equilibrated against 500 μL of reservoir solution containing 35% MPD. Crystals typically appeared within 24-96 h. Prior to data collection, crystals were transferred to cryoprotectant solution consisting of precipitant supplemented to 25% MPD, soaked for two minutes, and flash-frozen in liquid nitrogen. Diffraction data were collected using a Rigaku MicroMax-003 X-ray source equipped with a Dectris Pilatus 200K detector. Data were indexed, integrated, and scaled using HKL3000.^30^ Crystallographic statistics are provided in **Table S9,10**.

#### Structure Determination and Refinement

Initial phases for all structures were obtained by molecular replacement in Phaser^31^ using the *env2* riboswitch model derived from PDB 9E5I^21^. To minimize model bias, the ligand and nucleotide A20 were removed from the search model. Ligand coordinates were manually built in PyMol (v2.5.7, Schrödinger) from the previously determined structure of Cbl **4** and restraint files were prepared using eLBOW.^32^ Following initial refinement, ligands were placed into unambiguous electron density using LigandFit^33^ and A20 was subsequently modeled in Coot (0.9.8.93).^34^ Iterative cycles of model building and refinement were performed in PHENIX (1.19.2),^35^ including multiple rounds of maximum-likelihood refinement and simulated annealing in torsion space (5,000 K) to further reduce model bias.^36^ Solvent molecules were added where supported by electron density and RNA geometry was optimized using ERRASER.^37^ To validate placement of the ligand and A20, simulated-annealing 2F_o_-F_c_ omit maps were calculated for all structures in which both the ligand and A20 were excluded from the model prior to refinement.

#### Cheminformatic and Machine Learning (ML) Analysis

Molecular descriptors were calculated in MOE for the alkyne synthetic precursors corresponding to the variable β-axial substituents within the Cbl derivative library. Descriptor calculations were performed on energy-minimized structures as described above. A total of 153 two- and three-dimensional descriptors spanning size, shape, polarity, surface area, flexibility, topology, and electronic properties were retained for analysis. Natural log-transformed *env8* K_D_ values [ln(K_D_)] were used as the response variable for all modeling analyses.

Cheminformatic and ML analyses were performed in MATLAB (R2024b) using elastic net (EN), random forest (RF), and gradient boosting (GB) regression models. Model performance was evaluated using 100 repeated holdout iterations with stratified 80/20 train-test splits generated from affinity-based response bins to preserve the distribution of binding affinities across training and test sets. For EN models, descriptors were standardized using parameters derived from the training data and model hyperparameters were optimized by five-fold cross-validation across a range of mixing parameters (α = 0.1-1.0). Sparse models containing five or fewer descriptors were preferentially selected. RF and GB models were constructed using ensembles of 300 regression trees with a minimum leaf size of three observations.

Model performance was assessed using root-mean-square error (RMSE), coefficient of determination (R^2^), and Pearson correlation coefficients (*r*) calculated on held-out test data. Descriptor importance was evaluated independently using EN descriptor selection frequencies and RF feature importance rankings. Stable descriptors were defined as those selected in at least 30% of EN iterations and were subsequently used to construct a final consensus model. Differences in descriptor distributions among aromatic, hybrid, and aliphatic ligand classes were evaluated using Wilcoxon rank-sum tests and Cliff’s Δ effect sizes.

## Data availability

Atomic coordinates and structure factors have been deposited in the Protein Data Bank (PDB) (https://www.rcsb.org/) under accession numbers 37CM (*env2*-Cbl **67**), 37CN (*env2*-Cbl **69**), 37CO (*env2*-Cbl **71**), 37CP (*env2*-Cbl **68**), 37CQ (*env2*-Cbl **70**), 37CR (*env2*-Cbl **72**).

## Author Contributions

L.T.O. and R.T.B. designed the research. D.P., A.J.W., and L.T.O. carried out small molecule synthesis and characterization. L.T.O. collected and analyzed all experimental and computational data and wrote the paper with feedback from all authors. R.T.B. supervised the project.

## Notes

The authors declare the following competing financial interest(s): R.T.B. serves on the Scientific Advisory Boards of SomaLogic, MeiraGTx, and Mol Horizon. R.T.B. and L.T.O. hold equity stakes in Mol Horizon.

## Acknowledgements

We thank the Macromolecular X-ray Crystallography Core (RRID:SCR_019310) at the University of Colorado Boulder for crystallographic data collection. We acknowledge A. Erbse for her assistance with the resources utilized at the University of Colorado Boulder.

