## Supplemental Information for "Reimagining productive chemical space for RNA recognition beyond aromaticity"

**This file includes:**


| | |
|---|---|
| Tables S1-S16 | 2-17 |
| Figures S1-S6 | 18-23 |
| Derivative synthesis and characterization | 24-38 |
| References | 39 |

**Table S1.** RNA-binding repository accession information.

| Repository name[a] | Number of compounds | Reference |
|---|---|---|
| ROBIN[b] | 2,003 | 1 |
| InfoRNA[c] | 508 | 2 |
| R-BIND[d] | 159 | 3 |
| DRTL[e] | 31 | 4 |
| ChemDiv ("Diverse")[f] | 50,000 | |

a. For chemical space comparison using RNA-binding repositories, we used ROBIN,[1] InfoRNA,[2] R-BIND,[3] and DRTL[4] datasets, which host known RNA-binding small molecules.
b. ROBIN dataset was accessed from: https://figshare.com/articles/dataset/Machine_Learning_Informs_RNA-Binding_Chemical_Space/20401974?file=36477873.
c. InfoRNA dataset was graciously provided by Amirhossein Taghavi and Matt Disney.
d. R-BIND dataset was accessed from: http://rbind.chem.duke.edu/ (using the "SM" dataset).
e. DRTL dataset was manually curated from the small molecule binders that emerged from the Duke RNA-Targeted Library (DRTL) library.
f. To compare chemically diverse matter to the RNA-binding repositories, we used the ChemDiv "Diverse 50k" collection (https://www.chemdiv.com/catalog/diversity-libraries/representative-diversity-libraries-out-of-1-6m-stock/).

**Table S2.** RNA-focused library accession information.

| Chemical supplier[a] | Number of compounds |
|---|---|
| Life chemicals (LC)[b] | 29,000 |
| Enamine[c] | 28,000 |
| ChemDiv[d] | 24,000 |
| MedChemExpress (MCE)[e] | 5,544 |
| Otava[f] | 2,391 |
| Asinex[g] | 1,862 |

a. For chemical space comparison using RNA-focused libraries, we chose collections from six vendors: Life chemicals (LC), Enamine, ChemDiv, MedChemExpress (MCE), Otava, and Asinex. In instances where certain vendors had more than one RNA library, we selected the largest and most general set. Additional information on the specific libraries is included below. To compare chemically diverse matter to the RNA-binding libraries, we used the ChemDiv "Diverse 50k" collection (see Table S1).

b. Life Chemicals (LC): "RNA Screening Library merged file 29k cmpds" (https://lifechemicals.com/screening-libraries/targeted-and-focused-screening-libraries/rna-focused-library).

c. Enamine: "RNA Library" (https://enamine.net/compound-libraries/targeted-libraries/rna).

d. ChemDiv: "RNA-Binding Chemical Space: (https://www.chemdiv.com/catalog/focused-and-targeted-libraries/rna-binding-chemical-space-library/).

e. MedChemExpress (MCE): "RNA Binding Library" (https://www.medchemexpress.com/screening/rna-binding-library.html).

f. Otava: "RNA Targeted Library: (https://www.otavachemicals.com/products/targeted-libraries-and-focused-libraries/rna-binding).

g. Asinex: "RNA-Targeting Small Molecules" (https://www.asinex.com/rna-binding).

**Table S3.** Cheminformatic descriptor[a,b] matrix for RNA-binding repositories.

| | ROBIN | | InfoRNA | | R-BIND | | DRTL | | Diverse | |
|---|---|---|---|---|---|---|---|---|---|---|
| **Descriptor** | **mean** | **SD** | **mean** | **SD** | **mean** | **SD** | **mean** | **SD** | **mean** | **SD** |
| arorings | 2.24 | 1.00 | 2.60 | 1.69 | 2.91 | 1.26 | 3.13 | 1.20 | 2.07 | 0.90 |
| b_rotN | 4.93 | 2.43 | 7.90 | 6.30 | 6.43 | 4.91 | 3.42 | 1.50 | 5.69 | 2.34 |
| b_rotR | 0.19 | 0.09 | 0.21 | 0.13 | 0.19 | 0.12 | 0.12 | 0.05 | 0.21 | 0.08 |
| chiral | 0.50 | 1.03 | 3.58 | 6.13 | 0.40 | 0.96 | 0.19 | 0.60 | 0.46 | 0.77 |
| logP | 2.73 | 1.69 | 1.48 | 4.84 | 2.97 | 2.37 | 4.64 | 2.54 | 2.46 | 1.36 |
| relPSA | 0.12 | 0.05 | 0.18 | 0.11 | 0.15 | 0.08 | 0.09 | 0.08 | 0.12 | 0.03 |
| Weight | 331.79 | 74.75 | 464.97 | 202.05 | 404.94 | 123.60 | 346.66 | 59.52 | 356.13 | 71.20 |

a. To provide a concise representation of molecular properties relevant to RNA recognition, seven descriptors spanning major physicochemical dimensions were selected for analysis: aromatic ring count (aromaticity; arorings), number of rotatable bonds (conformational flexibility; b_rotN), fraction rotatable bonds (relative flexibility; b_rotR), chiral centers (three-dimensionality; chiral), lipophilicity (logP (o/w))), relative polar surface area (polarity; relPSA), and molecular weight (molecular size; Weight).
b. All descriptors were calculated and defined within MOE (v2022.02).

**Table S4.** Cliff's Δ analysis[a,b,c] of descriptor matrix for RNA-binding repositories.

| Descriptor | ROBIN | InfoRNA | R-BIND | DRTL |
|---|---|---|---|---|
| arorings | 0.10 | 0.22 | 0.39 | 0.51 |
| b_rotN | -0.19 | 0.16 | -0.06 | -0.58 |
| b_rotR | -0.10 | -0.06 | -0.19 | -0.65 |
| chiral | 0.00 | 0.17 | -0.11 | -0.22 |
| logP | 0.12 | -0.03 | 0.13 | 0.52 |
| relPSA | -0.11 | 0.33 | 0.11 | -0.21 |
| Weight | -0.24 | 0.41 | 0.21 | -0.10 |

a. Cliff's Δ correspond to RNA-binding repository X – chemical diversity set ("Diverse").
b. 1 ≤ Δ ≥ 1, where Δ = 1: all values in group $x_i$ (repository X) are larger than group $j_i$ (diversity set), Δ = 0: values in repository X and diversity set are stochastically equal, Δ =- 1: all values in repository X are smaller than the diversity set. Effect-sizes are typically considered: small |Δ| > 0.147, medium: |Δ| > 0.330, and large: |Δ| > 0.474.
c. Wilcoxon rank-sum statistics were calculated but given the large samples sizes among groups, the resulting p-values were almost always ~0. As such, the main comparative contrast was the Cliff's Δ effect sizes.

**Table S5.** Cheminformatic descriptor[a,b,c] matrix for RNA-focused libraries.

| | LC | | Enamine | | ChemDiv | | MCE | | Otava | | Asinex | |
|---|---|---|---|---|---|---|---|---|---|---|---|---|
| **Descriptor** | **mean** | **sd** | **mean** | **sd** | **mean** | **sd** | **mean** | **sd** | **mean** | **sd** | **mean** | **sd** |
| arorings | 2.44 | 0.95 | 2.20 | 0.89 | 2.52 | 0.82 | 2.44 | 0.89 | 2.47 | 0.96 | 2.16 | 0.69 |
| b_rotN | 5.42 | 2.33 | 4.92 | 1.81 | 5.69 | 1.90 | 5.76 | 2.14 | 5.22 | 2.44 | 5.29 | 1.61 |
| b_rotR | 0.19 | 0.07 | 0.20 | 0.07 | 0.21 | 0.07 | 0.20 | 0.07 | 0.19 | 0.08 | 0.18 | 0.06 |
| chiral | 0.20 | 0.52 | 0.73 | 0.80 | 0.28 | 0.54 | 0.51 | 0.82 | 0.29 | 0.70 | 0.77 | 0.97 |
| logP | 3.05 | 1.45 | 1.60 | 1.38 | 3.04 | 1.41 | 3.25 | 1.14 | 2.73 | 1.32 | 1.93 | 1.23 |
| relPSA | 0.11 | 0.03 | 0.14 | 0.04 | 0.13 | 0.04 | 0.10 | 0.03 | 0.12 | 0.04 | 0.13 | 0.04 |
| Weight | 363.64 | 74.33 | 321.63 | 56.53 | 369.59 | 46.47 | 375.22 | 53.72 | 342.18 | 80.97 | 368.94 | 43.70 |

a. To provide a concise representation of molecular properties relevant to RNA recognition, seven descriptors spanning major physicochemical dimensions were selected for analysis: aromatic ring count (aromaticity; arorings), number of rotatable bonds (conformational flexibility; b_rotN), fraction rotatable bonds (relative flexibility; b_rotR), chiral centers (three-dimensionality; chiral), lipophilicity (logP (o/w))), relative polar surface area (polarity; relPSA), and molecular weight (molecular size; Weight).

b. All descriptors were calculated and defined within MOE (v2022.02).

c. Given that the descriptor values for the chemically diverse set are provided in Table S3, they have been omitted here.

**Table S6.** Cliff's Δ analysis[a,b,c] of descriptor matrix for RNA-focused libraries.

| Descriptor | LC | Enamine | ChemDiv | MCE | Otava | Asinex |
|---|---|---|---|---|---|---|
| arorings | 0.21 | 0.08 | 0.28 | 0.22 | 0.23 | 0.06 |
| b_rotN | -0.08 | -0.20 | 0.00 | 0.02 | -0.11 | -0.10 |
| b_rotR | -0.11 | -0.07 | 0.02 | -0.02 | -0.10 | -0.20 |
| chiral | -0.18 | 0.21 | -0.11 | 0.05 | -0.13 | 0.18 |
| logP | 0.23 | -0.36 | 0.25 | 0.32 | 0.12 | -0.26 |
| relPSA | -0.20 | 0.32 | 0.07 | -0.39 | -0.06 | 0.11 |
| Weight | 0.03 | -0.32 | 0.06 | 0.12 | -0.11 | 0.08 |

a. Cliff's Δ correspond to RNA library X – chemical diversity set ("Diverse").
b. 1 ≤ Δ ≥ 1, where Δ = 1: all values in group $x_i$ (library X) are larger than group $j_i$ (diversity set), Δ = 0: values in library X and diversity set are stochastically equal, Δ =- 1: all values in library X are smaller than the diversity set. Effect-sizes are typically considered: small |Δ| > 0.147, medium: |Δ| > 0.330, and large: |Δ| > 0.474.
c. Wilcoxon rank-sum statistics were calculated but given the large samples sizes among groups, the resulting p-values were almost always ~0. As such, the main comparative contrast was the Cliff's Δ effect sizes.

**Table S7.** Full-length *env8* Cbl riboswitch $K_D$ values.

| **Ligand**[a] | **$K_D$ (nM)**[b] |
|---|---|
| **1**, MeCbl | 1.4 ± 0.8 |
| **3**, CNCbl | 1.9 ± 0.7 |
| **57** | 120 ± 80 |
| **67** | 40 ± 10 |
| **68** | 30 ± 20 |
| **69** | 30 ± 20 |
| **70** | 35 ± 8 |
| **71** | 25 ± 9 |
| **72** | 24 ± 8 |
| **73** | 110 ± 30 |

a. Data for our newly synthesized amide-based cyclic hydrocarbon derivatives are shown with native ligands (MeCbl and CNCbl) and the unsubstituted amide **57** as reference.
b. Values are reported as mean and SD (*n* = 3) with rounding based on experimental uncertainty.

**Table S8.** Ligand-induced fold repression values of *env8*-GFP reporter.

| **Ligand**[a] | **Fold repression**[a] |
|---|---|
| **1**, MeCbl | 9.1 ± 0.2 |
| **3**, CNCbl | 10 ± 1 |
| **57** | 6 ± 2 |
| **67** | 6 ± 1 |
| **68** | 6 ± 1 |
| **69** | 5 ± 1 |
| **70** | 4 ± 1 |
| **71** | 4 ± 1 |
| **72** | 5 ± 1 |
| **73** | 2.6 ± 0.3 |

a. Data for our newly synthesized amide-based cyclic hydrocarbon derivatives are shown with native ligands (MeCbl and CNCbl) and the unsubstituted amide **57** as reference.
b. Values are reported as mean and SD (*n* = 4) with rounding based on experimental uncertainty.

**Table S9.** Crystallographic statistics of *env2* bound with Cbl derivatives.

| | ***env2*-Cbl 67** | ***env2*-Cbl 68** | ***env2*-Cbl 69** |
|---|---|---|---|
| **Data collection** | | | |
| Space group | P 1 21 1 | P 1 21 1 | P 1 21 1 |
| Cell dimensions | | | |
| a, b, c (Å) | 40.6, 78.6, 79.7 | 40.7, 78.2, 79.5 | 42.7, 76.5, 73.6 |
| α, β, γ (°) | 90.0, 91.0 90.0 | 90.0, 91.1, 90.0 | 90.0, 90.9, 90.0 |
| Wavelength (Å) | 1.54178 | 1.54178 | 1.54178 |
| Resolution (Å) | 19.92-2.50 | 20.00-2.80 | 20.00-2.50 |
| | (2.59-2.50)[a] | (2.90-2.80) | (2.59-2.50) |
| $R_{pim}$[b] | 0.031 (0.091) | 0.083 (0.173) | 0.074 (0.185) |
| *I* / σ | 19.1 (4.5) | 8.1 (2.7) | 10.3 (2.4) |
| Completeness (%) | 93.7 (70.4) | 90.1 (72.7) | 94.9 (79.4) |
| Redundancy | 2.8 (2.6) | 1.9 (1.5) | 2.4 (1.7) |
| $CC_{1/2}$ | 0.99 (0.99) | 0.97 (0.93) | 0.97 (0.94) |
| **Refinement** | | | |
| Resolution (Å) | 19.92-2.51 | 19.78-2.78 | 19.72-2.99 |
| No. Reflections | 16034 | 11302 | 9127 |
| $R_{work}$ / $R_{free}$ (%) | 18.9/25.4 | 19.5/25.6 | 21.0/29.7 |
| No. of atoms | 3668 | 3584 | 3514 |
| RNA | 3262 | 3262 | 3262 |
| Ligand | 198 | 200 | 202 |
| Co-solutes | 28 | 28 | 28 |
| Cations | 19 | 19 | 19 |
| Water | 161 | 75 | 3 |
| B-factors (Å$^2$) | | | |
| RNA | 30.7 | 24.3 | 14.2 |
| Ligand | 25.5 | 19.2 | 9.8 |
| Co-solutes | 30.9 | 25.1 | 23.7 |
| Cations | 30.4 | 23.4 | 15.9 |
| Water | 26.2 | 19.9 | 13.4 |
| R.M.S. deviations | | | |
| Bond length (Å) | 0.010 | 0.010 | 0.010 |
| Bond angles (°) | 1.83 | 1.82 | 1.81 |

[a]Values in parentheses are for the highest resolution shell.
[b]$R_{pim}$: precision-indicating merging R factor

**Table S10.** Crystallographic statistics of *env2* bound with Cbl derivatives.

| | ***env2*-Cbl 70** | ***env2*-Cbl 71** | ***env2*-Cbl 72** |
|---|---|---|---|
| **Data collection** | | | |
| Space group | P 1 21 1 | P 1 21 1 | P 1 21 1 |
| Cell dimensions | | | |
| a, b, c (Å) | 41.2, 78.1, 79.2 | 41.3, 78.4, 80.3 | 40.7, 78.4, 79.0 |
| α, β, γ (°) | 90.0, 90.8, 90.0 | 90.0, 89.3, 90.0 | 90.0, 91.4, 90.0 |
| Wavelength (Å) | 1.54178 | 1.54178 | 1.54178 |
| Resolution (Å) | 20.00-2.60 | 20.00-2.70 | 20.00-2.60 |
| | (2.69-2.60)[a] | (2.80-2.70) | (2.69-2.60) |
| $R_{pim}$[b] | 0.064 (0.154) | 0.056 (0.206) | 0.048 (0.132) |
| $I$ / σ | 12.8 (3.2) | 10.8 (1.9) | 15.0 (3.6) |
| Completeness (%) | 92.8 (70.1) | 92.7 (74.0) | 89.5 (67.8) |
| Redundancy | 2.9 (2.6) | 2.5 (2.1) | 2.7 (2.5) |
| $CC_{1/2}$ | 0.98 (0.96) | 0.99 (0.96) | 0.98 (0.95) |
| **Refinement** | | | |
| Resolution (Å) | 19.99-2.62 | 19.98-2.68 | 19.81-2.58 |
| No. Reflections | 13693 | 13045 | 13854 |
| $R_{work}$ / $R_{free}$ (%) | 21.1/27.8 | 22.1/27.9 | 20.1/27.0 |
| No. of atoms | 3588 | 3567 | 3617 |
| RNA | 3262 | 3262 | 3262 |
| Ligand | 204 | 204 | 206 |
| Co-solutes | 28 | 28 | 28 |
| Cations | 19 | 19 | 19 |
| Water | 75 | 54 | 102 |
| B-factors (Å$^2$) | | | |
| RNA | 35.5 | 32.6 | 32.1 |
| Ligand | 28.1 | 27.5 | 26.0 |
| Co-solutes | 35.7 | 34.2 | 33.7 |
| Cations | 31.8 | 28.9 | 32.6 |
| Water | 28.2 | 26.0 | 26.4 |
| R.M.S. deviations | | | |
| Bond length (Å) | 0.009 | 0.010 | 0.009 |
| Bond angles (°) | 1.74 | 1.78 | 1.64 |

[a]Values in parentheses are for the highest resolution shell.
[b]$R_{pim}$: precision-indicating merging R factor

**Table S11.** Comparison of descriptor selection frequencies/feature scores.

| Descriptor[a] | EN selection frequency[b] | RF importance[c], mean | RF importance, sd | RF importance, normalized[d] |
|---|---|---|---|---|
| npr1 | 77 | 0.0118 | 0.0037 | 100.00 |
| balabanJ | 2 | 0.0088 | 0.0039 | 74.20 |
| DCASA | 70 | 0.0024 | 0.0014 | 20.24 |
| a_nN | 68 | 0.0007 | 0.0006 | 5.77 |
| AM1_HF | 52 | 0.0080 | 0.0041 | 67.80 |
| GCUT_SLOGP_1 | 0 | 0.0077 | 0.0030 | 64.91 |
| SlogP_VSA1 | 60 | 0.0017 | 0.0011 | 14.56 |
| AM1_LUMO | 18 | 0.0063 | 0.0029 | 53.33 |
| dipoleX | 50 | 0.0009 | 0.0006 | 7.28 |
| E_str | 50 | 0.0046 | 0.0028 | 39.23 |
| PEOE_RPC+ | 0 | 0.0057 | 0.0030 | 48.63 |
| SMR_VSA4 | 12 | 0.0055 | 0.0032 | 46.36 |
| PEOE_VSA-0 | 46 | 0.0027 | 0.0017 | 22.67 |
| opr_brigid | 40 | 0.0012 | 0.0010 | 10.15 |

a. All descriptors were calculated and defined within MOE (v2022.02). Physical meaning and plain English explanation of each descriptor is found in Table S12.
b. Elastic net (EN) selection frequencies are defined as the number of times a given descriptor was selected by the model over 100 repeated cross-validation runs.
c. Random forest (RF) feature importance scores are defined as mean increase in prediction error resulting from permutation of each descriptor, averaged across the out-of-bag (OOB) samples of all trees.
d. Normalized RF feature importance scores scale the highest-scoring descriptor to 100.

**Table S12.** Consensus model-selected descriptors from RNA-ligand binding data.

| Descriptor[a] | Descriptor meaning | Plain English |
|---|---|---|
| npr1 | Normalized principal moment ratio | Molecular shape; distinguishes rod-like from compact molecular geometries |
| balabanJ | Balaban topological index | Overall molecular connectivity and branching complexity |
| DCASA | Difference in charge-weighted accessible surface area | Balance and distribution of charged versus neutral molecular surface |
| a_nN | Number of nitrogen atoms | Nitrogen (heteroatom) content available for hydrogen bonding and electrostatic interactions |
| AM1_HF | AM1 heat of formation | Overall molecular electronic stability and energetic character |
| GCUT_SLOGP_1 | Graph-based logP descriptor | Combined measure of molecular topology and lipophilicity |
| SlogP_VSA1 | Surface area within a defined logP range | Surface area associated with weakly lipophilic regions of the molecule |
| AM1_LUMO | Lowest unoccupied molecular orbital (LUMO) energy | Ability to accept electron density; molecular electrophilicity |
| dipoleX | X-component of molecular dipole moment | Directional molecular polarity along the principal X-axis |
| E_str | Bond stretching energy | Internal strain associated with bond deformation |
| PEOE_RPC+ | Relative positive partial charge | Magnitude and distribution of positive electrostatic charge |
| SMR_VSA4 | Surface area within a defined molar refractivity range | Surface area associated with regions of intermediate polarizability |
| PEOE_VSA-0 | Surface area with weakly negative partial charge | Accessible molecular surface carrying small negative electrostatic charge |
| opr_brigid | Rigid bond count | Degree of conformational rigidity within the molecular scaffold |

a. All descriptors were calculated and defined within MOE (v2022.02).

**Table S13.** Coefficients of the final consensus elastic net regression model.

| Term | Estimate[a] | p-value[b] |
|---|---|---|
| Intercept | -16.78 | 1.43E-85 |
| AM1_HF | -0.35 | 0.01 |
| a_nN | 0.10 | 0.66 |
| DCASA | -0.29 | 0.04 |
| dipoleX | -0.12 | 0.41 |
| npr1 | 0.44 | 1.95E-4 |
| SlogP_VSA1 | -0.27 | 0.14 |

a. Estimate of the regression coefficient for a given descriptor, holding the other descriptors constant.
b. The p-value tests the null hypothesis that the regression coefficient for a given descriptor is equal to zero in the final multiple linear regression model.

**Table S14.** Cliff's Δ analysis[a] of descriptor distributions among ligand chemotypes.

| **Descriptor[b]** | **Mean (aromatic, hybrid, aliphatic)** | **Aromatic vs hybrid (Δ, p-value)[c]** | **Aromatic vs aliphatic (Δ, p-value)[d]** | **Hybrid vs aliphatic (Δ, p-value)[e]** |
|---|---|---|---|---|
| npr1 | 0.20, 0.17, 0.19 | 0.09, 0.75 | -0.01, 0.96 | -0.10, 0.75 |
| DCASA | 118.98, 72.94, 116.67 | 0.10, 0.70 | -0.03, 0.87 | -0.14, 0.65 |
| a_nN | 0.45, 0.33, 0.65 | -0.13, 0.47 | -0.41, 0.002 | -0.31, 0.21 |
| SlogP_VSA1 | 4.51, 7.34, 6.28 | -0.38, 0.06 | -0.54, 0.0003 | 0.06, 0.85 |
| AM1_HF | 66.22, 44.08, -7.35 | 0.35, 0.17 | 0.67, <0.0001 | 0.80, 0.005 |
| dipoleX | 0.25, -0.04, 0.05 | 0.33, 0.20 | 0.25, 0.12 | -0.10, 0.75 |
| E_str | 0.54, 0.44, 0.24 | -0.13, 0.60 | 0.62, 0.0002 | 0.78, 0.006 |
| PEOE_VSA-0 | 34.69, 41.43, 30.41 | -0.23, 0.38 | 0.20, 0.23 | 0.47, 0.10 |
| opr_brigid | 8.64, 10.00, 3.29 | -0.31, 0.17 | 0.85, <0.0001 | 0.94, 0.0006 |
| AM1_LUMO | -0.25, -0.20, 0.75 | -0.18, 0.47 | -0.83, <0.0001 | -1.00, 0.0004 |

a. b. 1 ≤ Δ ≥ 1, where Δ = 1: all values in group $x_i$ are larger than group $j_i$, Δ = 0: values in $x_i$ and $j_i$ are stochastically equal, Δ =- 1: all values in $x_i$ are smaller than $j_i$ Effect-sizes are typically considered: small |Δ| > 0.147, medium: |Δ| > 0.330, and large: |Δ| > 0.474. Wilcoxon rank-sum p-values were also calculated.

b. All descriptors were calculated and defined within MOE (v2022.02). This set represents the 10 features most frequently selected by the elastic net model over 100 repeated cross-validation runs.

c. Cliff's Δ (and p-value calculations) correspond to aromatic – hybrid.

d. Cliff's Δ (and p-value calculations) correspond to aromatic – aliphatic.

e. Cliff's Δ (and p-value calculations) correspond to hybrid – aliphatic.

**Table S15.** RNA constructs used in this study.

| Name | Sequence (5′-to-3′) |
|---|---|
| *env8*_full length | GGC CUA AAA GCG UAG UGG GAA AGU GAC GUG AAA UUC GUC CAG AUU ACU UGA UAC GGU UAU ACU CCG AAU GCC ACC UAG GCC AUA CAA CGA GCA AGG AGA CUC A |
| *env2*_crystal construct | GGU AAA AGC GUA GUG GGA AAG UGA CGU GAA AUU CGU CCA GAC GAA AGU ACG GUU AUA CUC CGA AUG CCA CCU ACC A |

**Table S16.** List of alkynes used to synthesize Cbls **67-73**.

| Name | Desired Cbl | CAS # | Product #[a] |
|---|---|---|---|
| N-cyclopropylprop-2-ynamide | **67** | 1207294-09-4 | EN300-1237931 |
| N-(cyclopropylmethyl)prop-2-ynamide | **68** | 1849390-23-3 | EN300-1665582 |
| N-cyclopentylprop-2-ynamide | **69** | 1207294-10-7 | EN300-1265819 |
| N-(cyclopentylmethyl)prop-2-ynamide | **70** | 1855518-62-5 | EN300-2992778 |
| N-cyclohexylprop-2-ynamide | **71** | 22231-08-9 | EN300-6808733 |
| N-(cyclohexylmethyl)prop-2-ynamide | **72** | 1878553-75-3 | EN300-3013360 |
| N-benzylprop-2-ynamide | **73** | 87605-11-6 | EN300-306827 |

a. All alkynes were purchased from Enamine (https://enaminestore.com).

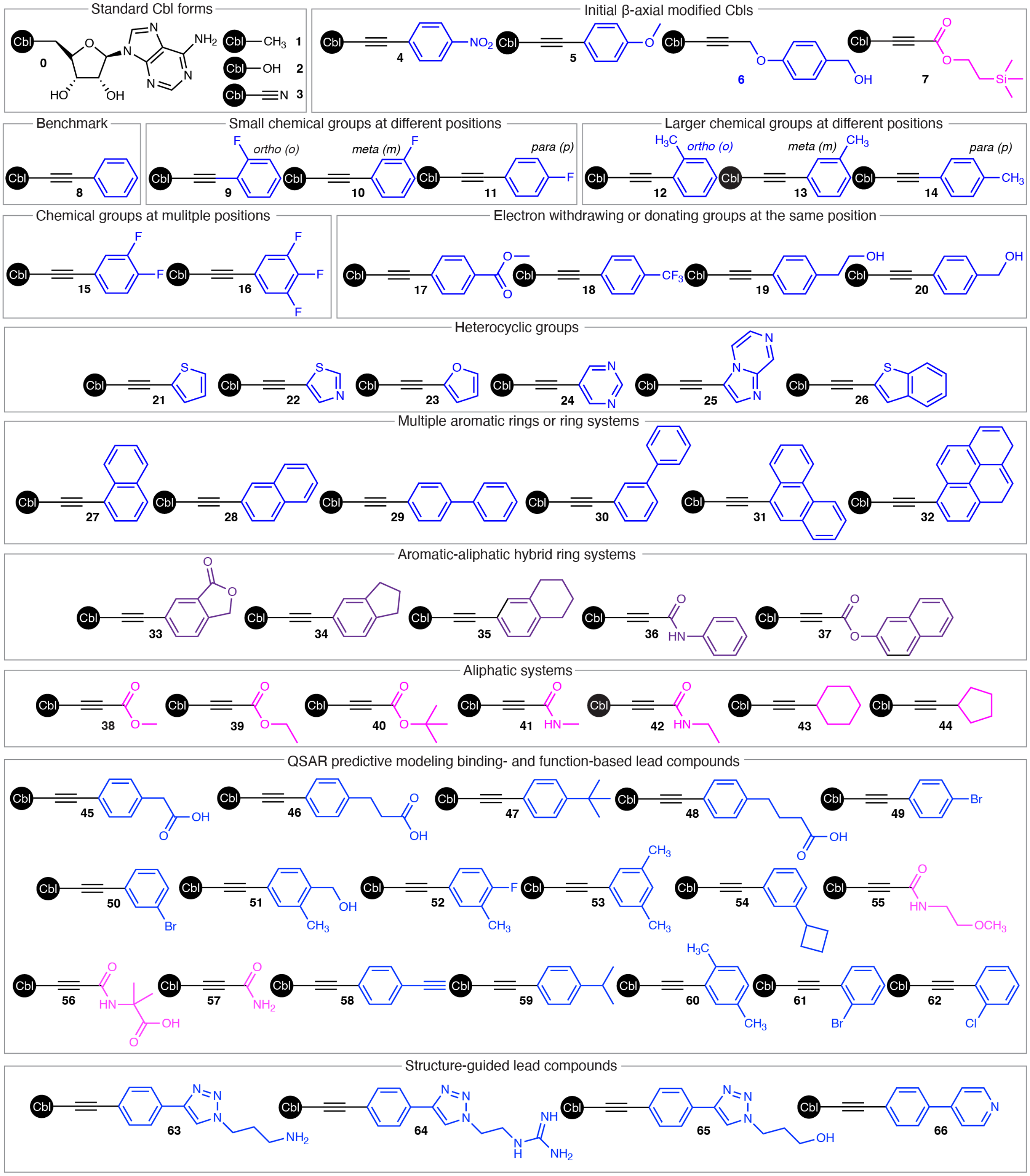


**Figure S1.** Overview of the Cobalamin (Cbl) β-axial derivative library. Structures of previously reported Cbl β-axial derivatives spanning aromatic, aliphatic, and aromatic-aliphatic hybrid chemotypes. Ligands are grouped according to the design strategy used during library development. Native Cbl forms and the initial β-axial modified derivatives used to establish the platform are shown for reference.

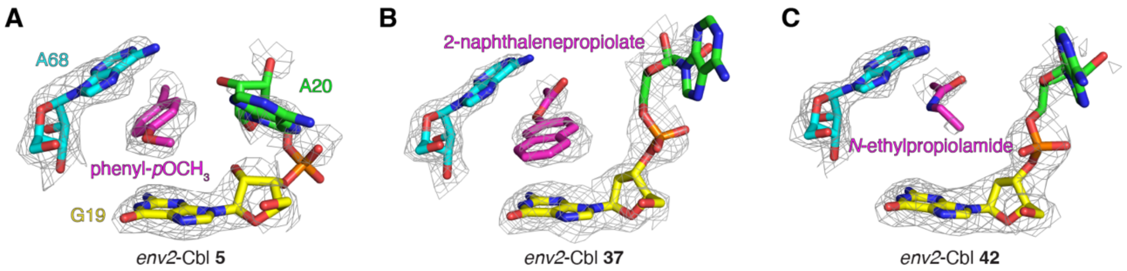


**Figure S2.** Representative recognition modes among diverse chemotypes within the *env2* cryptic binding site. Crystal structures of aromatic phenyl-*p*$OCH_3$-Cbl (**5**) (A), aromatic-aliphatic hybrid 2-naphthalenepropiolate-Cbl (37) (B), and aliphatic *N*-ethylpropiolamide-Cbl (**42**) (C) bound to the homologous *env2* cobalamin riboswitch aptamer domain. Binding-pocket nucleotides G19 (yellow), A20 (green), and A68 (cyan) are numbered according to the full-length *env8* riboswitch and colored accordingly; β-axial substituents are shown in magenta. Mesh representations correspond to simulated-annealing $2F_o$-$F_c$ omit maps in which A20 and the ligand were omitted from the model and are contoured at 1σ.

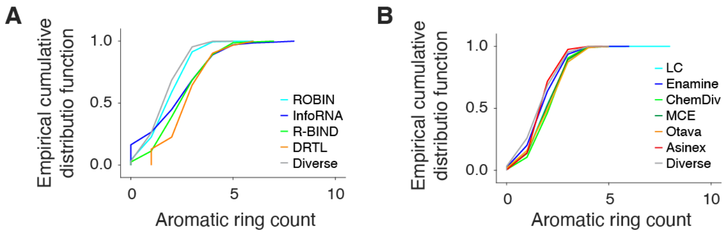


**Figure S3.** Empirical cumulative distribution function (ECDF) analysis of aromatic ring counts. (A) ECDF plots showing the distribution of aromatic ring counts among publicly available RNA-binding ligand repositories (ROBIN, Inforna, R-BIND, and DRTL) relative to a chemically diverse reference collection from ChemDiv (“Diverse”). (B) ECDF plots showing the distribution of aromatic ring counts among commercially available RNA-focused screening collections, including Life Chemicals (LC), Enamine, ChemDiv, MedChemExpress (MCE), Otava, and Asinex, in reference to the same “Diverse” set.

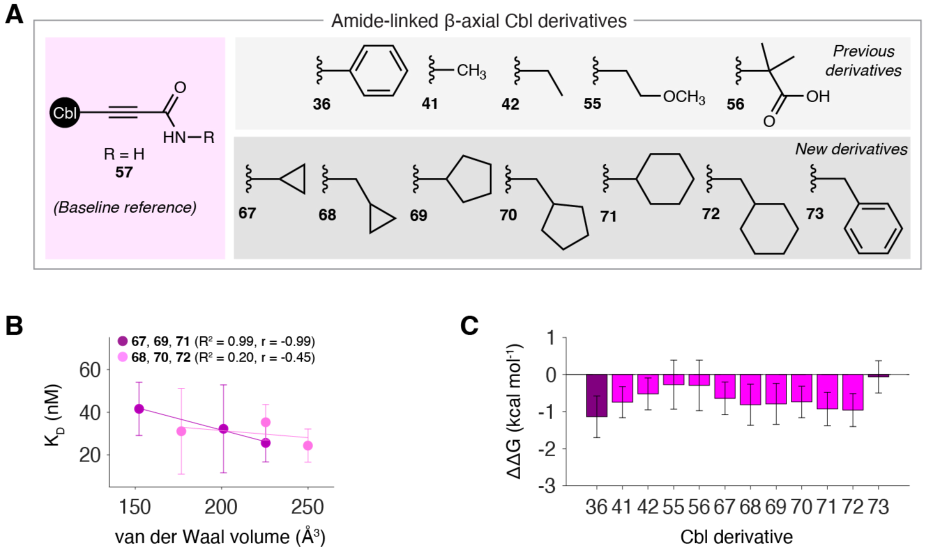


**Figure S4.** Structure-activity relationships (SAR) within the amide-linked β-axial Cbl derivative series. (A) Structures of previously reported amide-linked β-axial derivatives (**36**, **41**, **42**, **55**, and **56**), the unsubstituted amide reference compound **57**, and newly synthesized derivatives **67-73**. (B) SAR between RNA-binding affinity ($K_D$) and calculated van der Waals volume for the cyclic hydrocarbon series. Dark and light magenta data sets correspond to directly linked (**67**, **69**, and **71**) and spacer-containing (**68**, **70**, and **72**) derivatives, respectively. Linear regression statistics are shown. (C) Relative binding free energies (ΔΔG) for amide-linked β-axial derivatives calculated relative to the unsubstituted amide reference compound **57**. Error bars represent propagated uncertainty from independent binding measurements.

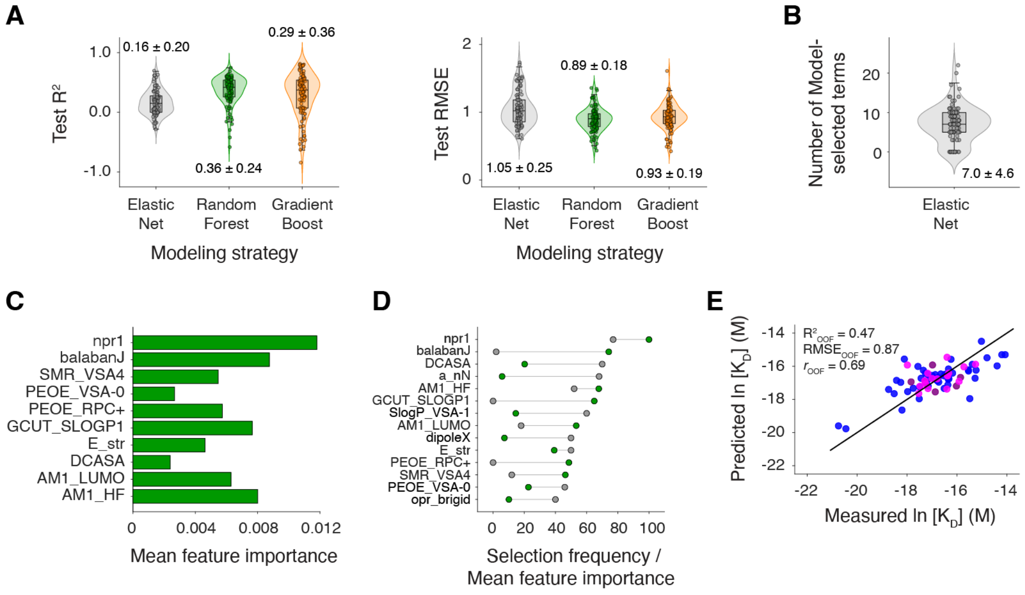


**Figure S5**. Additional machine learning model benchmarking and descriptor-selection analyses. (A) Distributions of test-set $R^2$ values (left) and root-mean-square error (RMSE) (right) obtained from 100 repeated train-test iterations using elastic net (EN), random forest (RF), and gradient boosting (GB) regression models. Values indicate mean ± SD. (B) Distribution of the number of descriptors selected by EN models across all modeling iterations. Values indicate mean ± SD. (C) Mean RF feature importance scores for the ten highest-ranked descriptors. (D) Comparison of EN descriptor selection frequencies (gray) and RF feature importance rankings (green) for descriptors identified as important by one or both modeling approaches. Here, RF feature importance is normalized so that the highest-scoring descriptor is scaled to 100. (E) Out-of-fold (OOF) predictions from the GB model. Model performance statistics are shown.

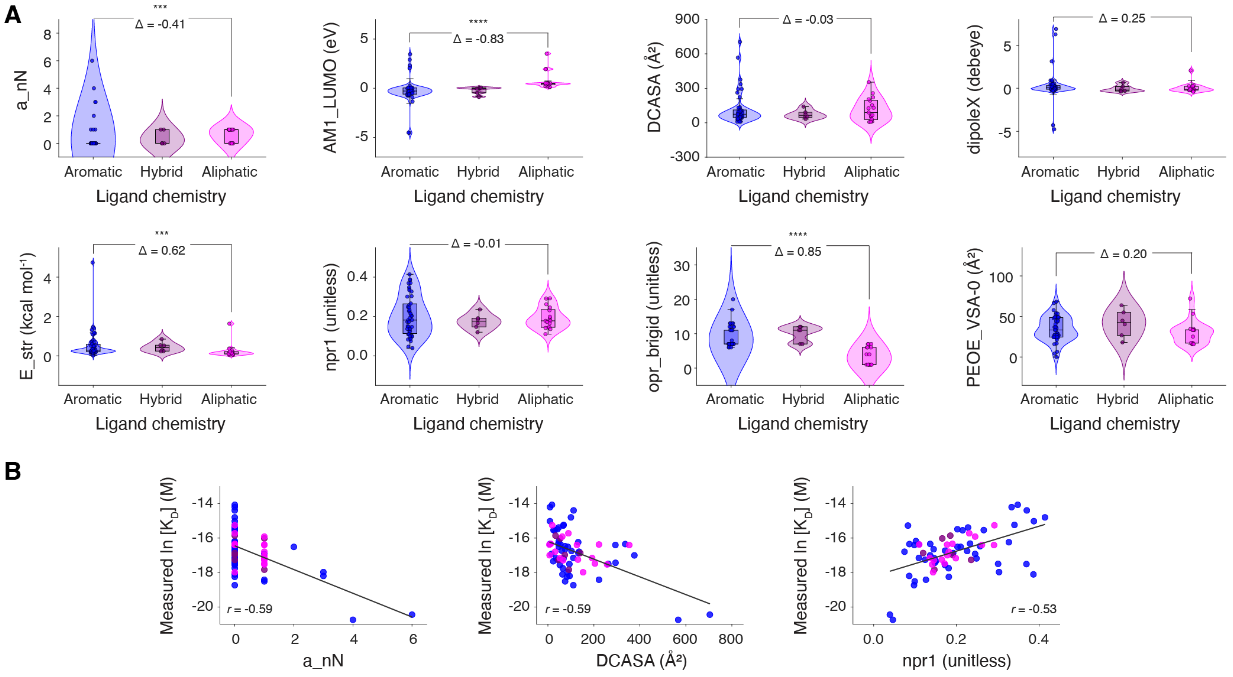


**Figure S6.** Additional descriptor analyses associated with *env8* RNA binding. (A) Distributions of descriptor values across aromatic, hybrid, and aliphatic ligand classes for descriptors identified as important by EN and/or RF modeling approaches. Statistical significance was assessed using Wilcoxon rank-sum tests (***, $p < 0.001$; ****, $p < 0.0001$) and Cliff's Δ effect size. (B) Relationships between measured binding affinity (natural-log transformed $K_D$) and descriptors a_nN, DCASA, and npr1. Pearson correlation coefficients (*r*) are shown.

### Derivative synthesis and characterization

#### General

Commercially available reagents and solvents were used as received. $^1$H NMR spectra were recorded at room temperature on a Bruker 400 MHz spectrometer with the residual solvent peak used as an internal standard. Data are reported as follows: chemical shift, peak multiplicity (s = singlet, d = doublet, t = triplet, q = quartet, m = multiplet), coupling constants (Hz), and number of protons. For clarity, integration values are shown only for selected resonances in the spectra. Due to restricted rotation on the NMR timescale, selected resonances are observed as two signals corresponding to rotamers present in an approximately 1:1 ratio. For these resonances, signals corresponding to a single proton are reported with an integral of 0.5 H for each rotamer. HPLC and HRMS analyses were performed to confirm the identity and purity of all derivatives. HRMS were recorded on Waters Cyclic IMS qTOF. All reactions and product purities were monitored using RP-HPLC techniques. HPLC analytical measurement conditions: column, Phenomenex Jupiter 5 µm C18 300 Å, 250 mm, 4.6 mm; detection, UV/Vis (λ = 254 and 361 nm); pressure, 10 MPa; temperature, 22 ºC; flow, 1 mL/min; gradient, 10% MeCN/$H_2O$ + 0.02% TFA to 70% MeCN/$H_2O$ + 0.02% TFA in 16 min. Abbreviations: DMA, dimethylacetamide; DBU, 1,8-Diazabicyclo[5.4.0]undec-7-ene; $Cu(OAc)_2$, copper(II) acetate; DMSO, dimethyl sulfoxide; $Et_2O$, diethyl ether; MeCN, acetonitrile; MeOH, methanol; RP-HPLC, reverse-phase high-performance liquid chromatography; TFA, trifluoroacetic acid; HRMS, high-resolution mass spectroscopy; Da, dalton; $t_R$, retention time.

#### General procedure for synthesis of β-axial-modified Cbls 67-73

β-axial-modified Cbl derivatives were synthesized according to our previously reported methodology.[5–7] Briefly, CNCbl (30 mg, 0.022 mmol) was dissolved in DMA, followed by addition of the appropriate alkyne (0.22 mmol; Table S16), $Cu(OAc)_2$ (2.7 mg, 0.022 mmol), and DBU (6.7 µL, 0.044 mmol). The reaction mixture was stirred vigorously at room temperature for 3 h, with progress monitored by HPLC. Upon near-complete consumption of the starting material, MeOH (3 mL) and $Et_2O$ (30 mL) were added sequentially to precipitate the product. The resulting solid was collected by centrifugation, air-dried, and redissolved in $H_2O$ (5 mL). Following centrifugation to remove insoluble copper-containing species, the supernatant was purified by reverse-phase C18 chromatography using a gradient of $H_2O$/MeCN (10-25% MeCN). Fractions containing the desired product, as confirmed by HPLC, were combined and concentrated under reduced pressure. The residue was dissolved in MeOH (1 mL), reprecipitated with $Et_2O$ (15 mL), collected by centrifugation, and dried under reduced pressure overnight to afford the corresponding β-axial Cbl derivative.

### Characterization of Cbl derivatives

Cbl **67**: Yield: 16.2 mg (51%); HR HRMS (ESI) m/z $[M + H]^+$ calcd for $C_{68}H_{95}CoN_{14}O_{15}P^+$, 1,427.6165 Da; found, 1,437.6147 Da; $t_R$ (RP-HPLC): 8.59 min; $^1H$ NMR (400 MHz, DMSO-$d_6$): δ 7.73 – 7.47 (m, 5H), 7.39 – 7.25 (m, 3H), 7.18 (s, 1H), 7.12 (s, 1H), 7.06 (s, 0.5H), 7.03 (s, 0.5H), 6.99 (d, *J* = 9.4 Hz, 1H), 6.94 (s, 1H), 6.80 (s, 1H), 6.67 (s, 1H), 6.55 (s, 1H), 6.49 – 6.38 (m, 2H), 6.26 (s, 1H), 6.11 – 6.04 (m, 1H), 5.85 (s, 1H), 5.81 (s, 1H), 4.47 (t, *J* = 11.3 Hz, 1H), 4.26 (d, *J* = 8.3 Hz, 1H), 4.17 – 4.08 (m, 2H), 3.91 – 3.82 (m, 2H), 3.78 – 3.69 (m, 1H), 3.60 – 3.48 (m, 3H), 3.07 (d, *J* = 10.4 Hz, 1H), 2.73 – 2.63 (m, 3H), 2.46 – 2.35 (m, 10H), 2.32 – 2.21 (m, 3H), 2.21 – 2.11 (m, 6H), 2.10 – 1.49 (m, 12H), 1.66 (s, 3H), 1.56 – 1.51 (m, 1H), 1.33-1.30 (m, 3H), 1.28 – 1.13 (m, 8H), 1.10 – 1.02 (m, 4H), 0.98 (s, 1H), 0.96 – 0.88 (m, 1H), 0.50 – 0.45 (m, 1H), 0.34 – 0.19 (m, 5H).

HPLC chromatogram (λ = 254 and 361nm):

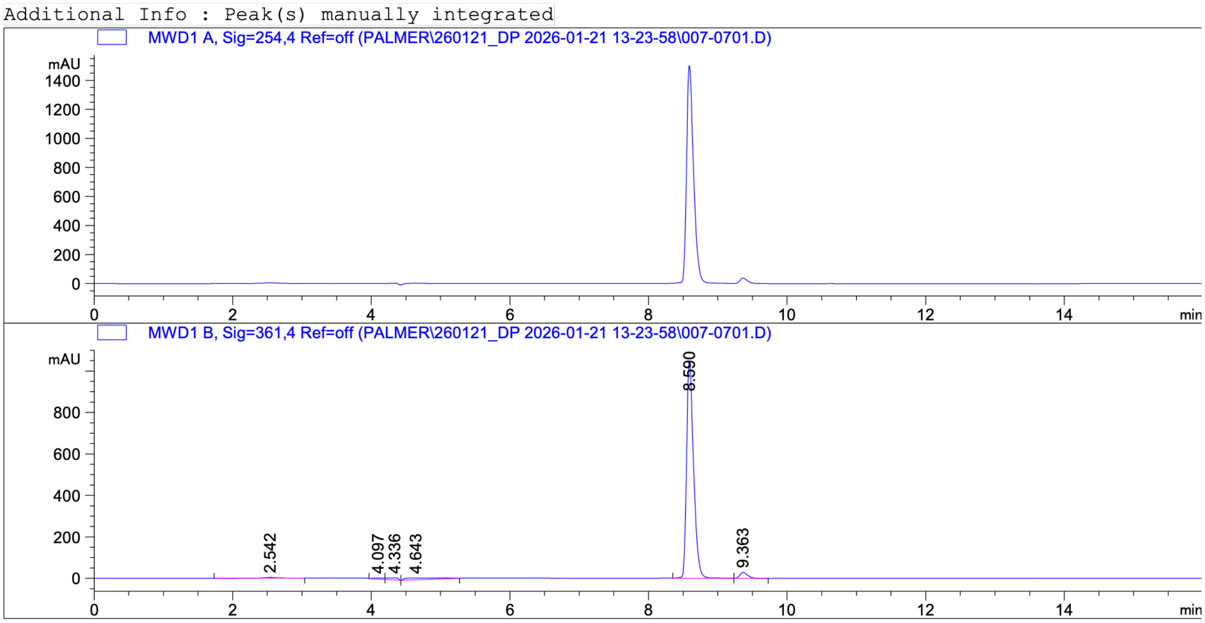

### HRMS spectrum:

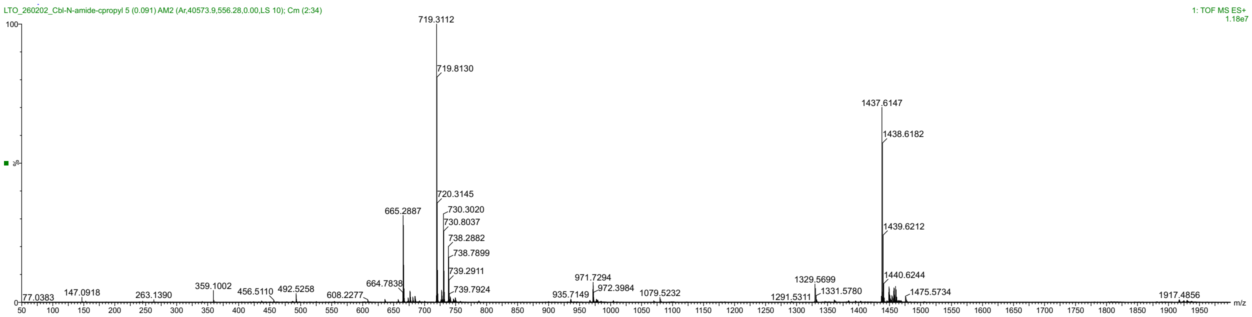


### [1]H NMR spectrum (400 MHz, DMSO-d$_6$):

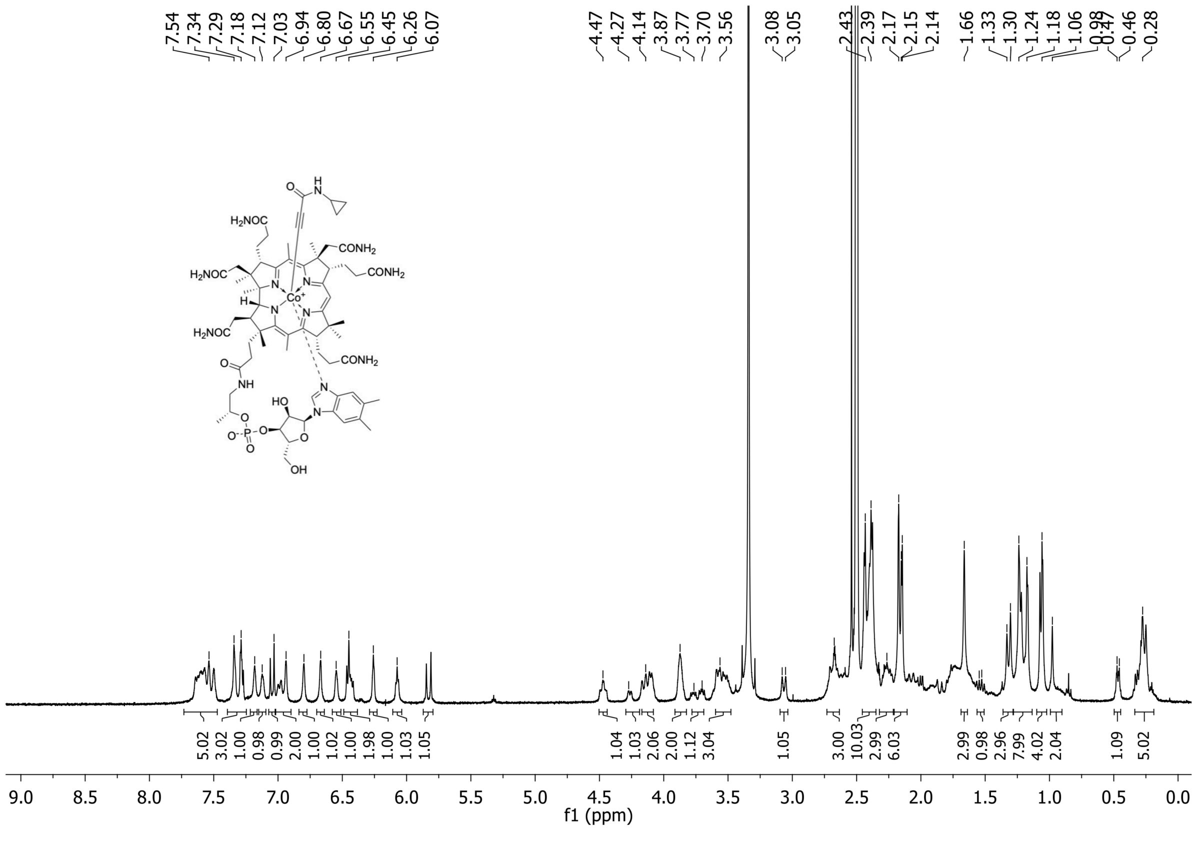

Cbl **68**: Yield: 18.9 mg (59%); HRMS (ESI) m/z $[M + H]^+$ calcd for $C_{69}H_{97}CoN_{14}O_{15}P^+$, 1,451.6322 Da; found, 1,451.6301 Da; $t_R$ (RP-HPLC): 8.97 min; $^1H$ NMR (400 MHz, DMSO-$d_6$): δ 7.71 – 7.49 (m, 5H), 7.35 – 7.33 (m, 1H), 7.29 (s, 1H), 7.18 – 7.15 (m, 1H), 7.11 (s, 1H), 7.05 (s, 0.5H), 7.04 (s, 0.5H), 7.00 (s, 0.5H), 6.97 (s, 0.5H), 6.94 (s, 1H), 6.80 (s, 1H), 6.68 (s, 0.5H), 6.66 (s, 0.5H), 6.55 (s, 0.5H), 6.54 (s, 0.5H), 6.45 (s, 1H), 6.44 – 6.38 (m, 1H), 6.26 (s, 1H), 6.08 (s, 1H), 5.85 (s, 0.5H), 5.83 (s, 0.5H), 4.47 (td, *J* = 9.0, 4.0 Hz, 1H), 4.18 – 4.05 (m, 3H), 3.92 – 3.83 (m, 2H), 3.78 – 3.69 (m, 1H), 3.61 – 3.49 (m, 3H), 3.07 (d, *J* = 11.2 Hz, 1H), 2.76 – 2.62 (m, 5H), 2.62 – 1.69 (m, 12H), 2.44 – 2.38 (m, 10H), 2.33 – 2.21 (m, 3H), 2.17 (s, 3H), 2.15 (s, 3H), 1.66 (s, 3H), 1.57 – 1.50 (m, 1H), 1.33 (d, *J* = 8.4 Hz, 3H), 1.23 (s, 3H), 1.17 (d, *J* = 7.6 Hz, 3H), 1.06 (d, *J* = 4.4 Hz, 4H), 1.02 (s, 1H), 0.95 – 0.85 (m, 1H), 0.81 – 0.68 (m, 1H), 0.67 – 0.50 (m, 1H), 0.36 – 0.18 (m, 5H), 0.04 – -0.02 (m, 1H), -0.04 – -0.11 (m, 1H).

HPLC chromatogram (λ = 254 and 361nm):

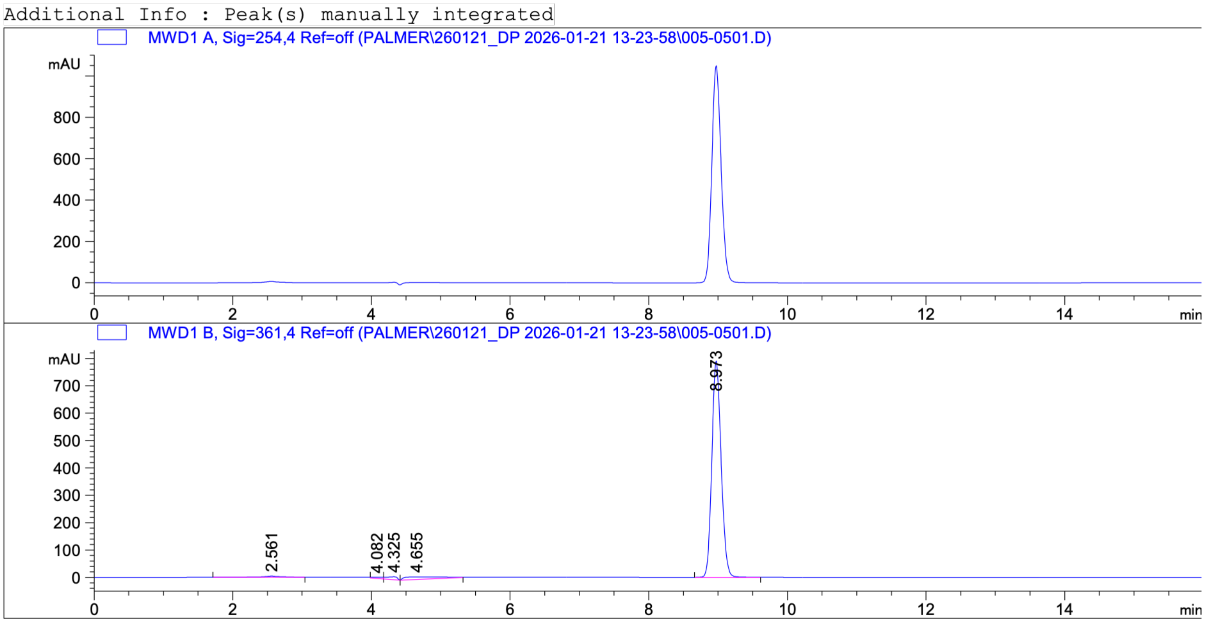

### HRMS spectrum:

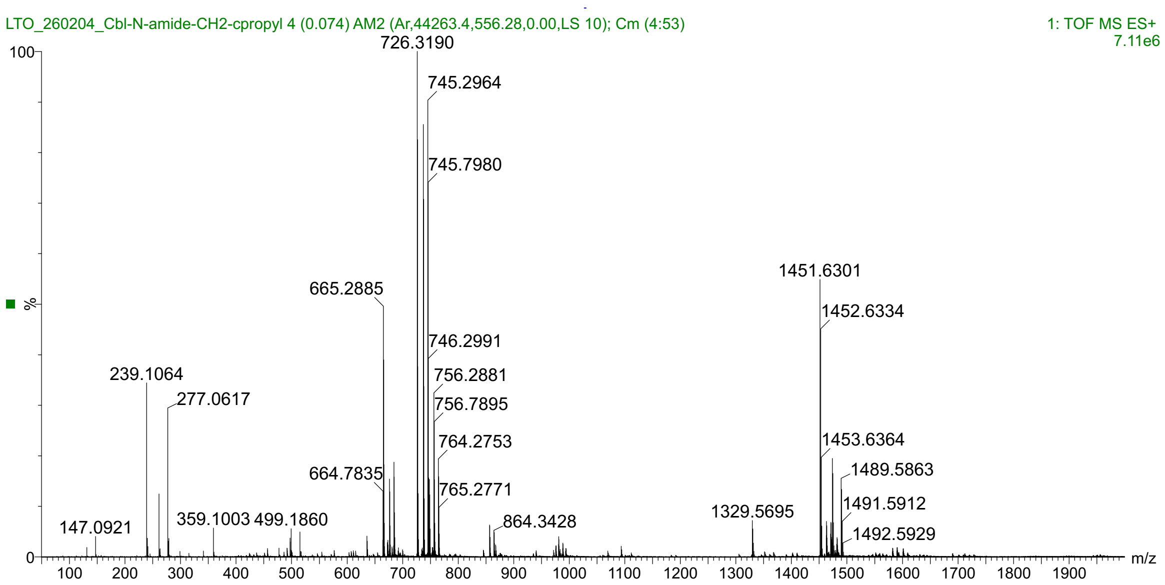


### $^{1}H$ NMR spectrum (400 MHz, DMSO-$d_6$):

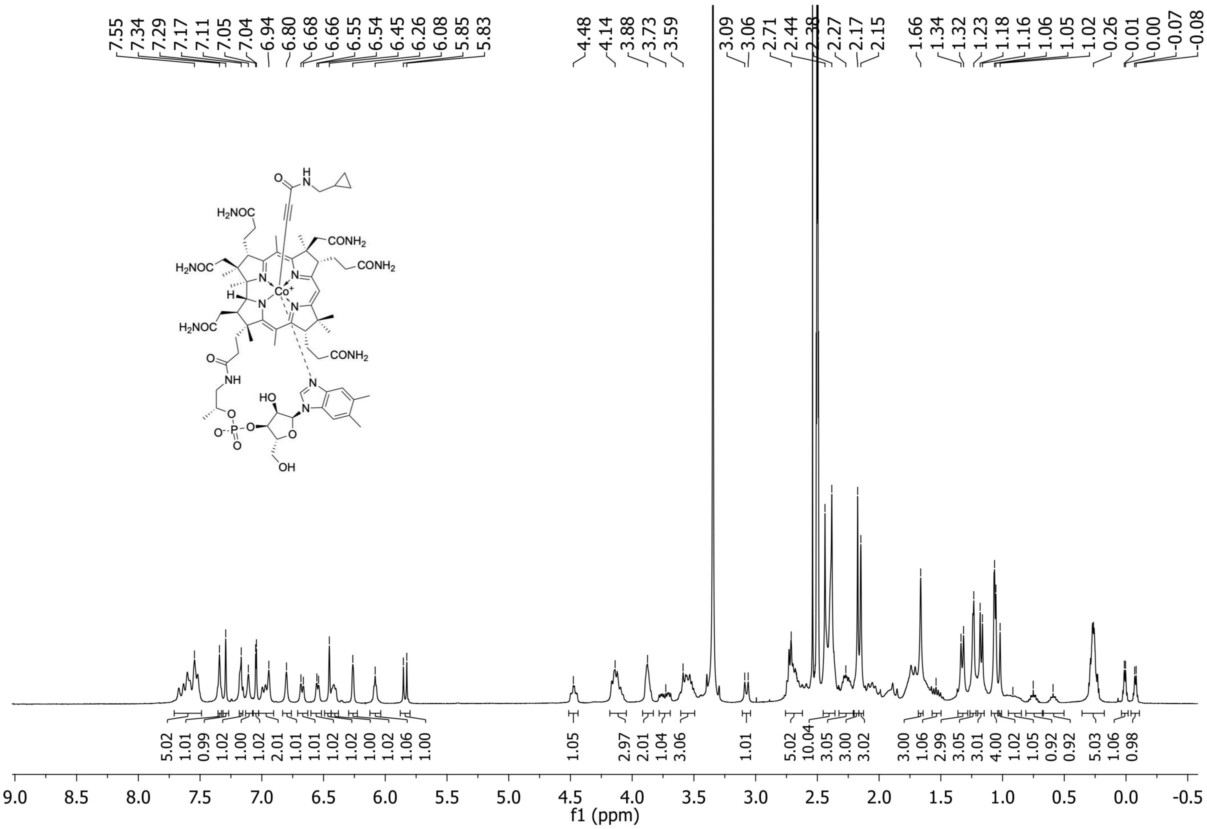

Cbl **69**: Yield: 15.5 mg (48%); HRMS (ESI) m/z $[M + H]^+$ calcd for $C_{70}H_{99}CoN_{14}O_{15}P^+$, 1,465.6478 Da; found, 1,465.6456 Da; $t_R$ (RP-HPLC): 9.35 min; $^1H$ NMR (400 MHz, DMSO-$d_6$): δ 7.75 – 7.44 (m, 5H), 7.34 (s, 1H), 7.29 (s, 1H), 7.17 (s, 1H), 7.12 (s, 1H), 7.05 (s, 0.5H), 7.04 (s, 0.5H), 7.03 – 6.90 (m, 2H), 6.80 (s, 1H), 6.67 (s, 1H), 6.56 (s,0.5H), 6.54 (s, 0.5H), 6.45 (s, 1H), 6.44 – 6.39 (m, 1H), 6.26 (d, *J* = 2.1 Hz, 1H), 6.08 (d, *J* = 3.8 Hz, 1H), 5.86 (s, 0.5H), 5.82 (s, 0.5H), 4.47 (td, *J* = 8.5, 3.4 Hz, 1H), 4.24 – 4.04 (m, 3H), 3.93 – 3.82 (m, 2H), 3.81 – 3.49 (m, 5H), 3.07 (d, *J* = 11.3 Hz, 1H), 2.74 – 2.60 (m, 3H), 2.47 – 2.32 (m, 12H), 2.32 – 2.21 (m, 3H), 2.20 – 2.13 (m, 6H), 2.12 – 1.55 (m, 11H), 1.66 (s, 3H), 1.54 – 1.44 (m, 4H), 1.40 – 1.10 (m, 4H), 1.32 (d, *J* = 15.0 Hz, 3H), 1.24 (s, 3H), 1.17 (s, 3H), 1.07 (s, 3H), 1.06 (s, 1H), 1.00 (s, 2H), 0.92 – 0.84 (m, 1H), 0.27 (s, 1.5H), 0.25 (s, 1.5H).

HPLC chromatogram (λ = 254 and 361nm):

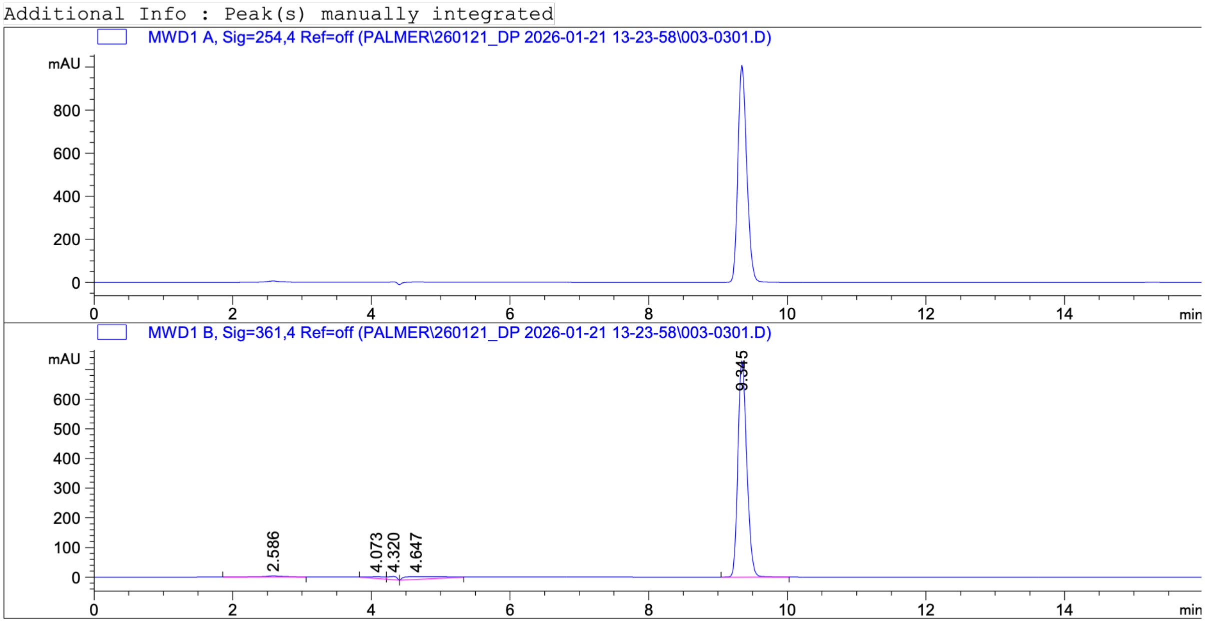

### HRMS spectrum:

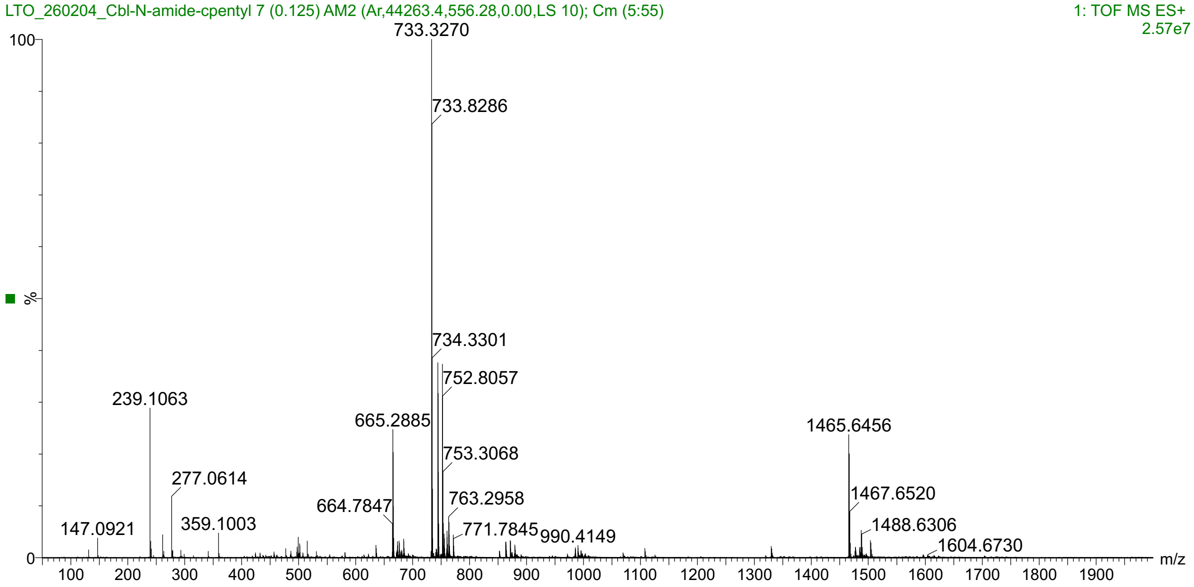


### $^{1}H$ NMR spectrum (400 MHz, DMSO-$d_6$):

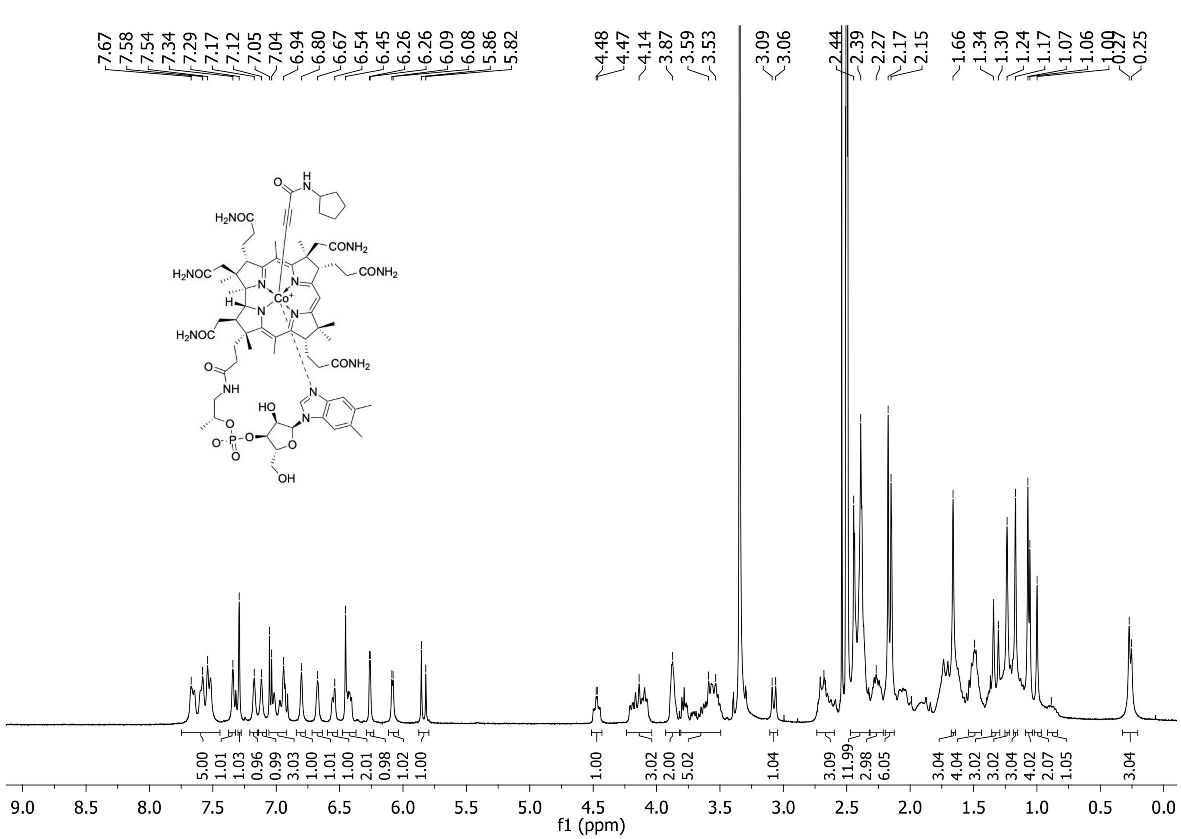

Cbl **70**: Yield: 18.1 mg (56%); HRMS (ESI) m/z [M + H]$^+$ calcd for $C_{71}H_{101}CoN_{14}O_{15}P^+$, 1,479.6635 Da; found, 1,479.6641 Da; $t_R$ (RP-HPLC): 9.80 min; $^1$H NMR (400 MHz, DMSO-$d_6$): δ 7.72 – 7.47 (m, 5H), 7.34 (s, 1H), 7.29 (s, 1H), 7.17 (s, 1H), 7.11 (s, 1H), 7.04 (s, 1H), 7.04 – 6.93 (m, 2H), 6.80 (s, 1H), 6.69 (s, 0.5H), 6.65 (s, 0.5H), 6.56 (s, 0.5H), 6.53 (s, 0.5H), 6.48 – 6.38 (m, 2H), 6.26 (s, 1H), 6.08 (s, 1H), 5.86 (s, 0.5H), 5.82 (s, 0.5H), 4.47 (td, *J* = 9.1, 3.1 Hz, 1H), 4.21 – 4.03 (m, 3H), 3.91 – 3.66 (m, 3H), 3.62 – 3.46 (m, 3H), 3.07 (d, *J* = 11.4 Hz, 1H), 2.80 – 2.60 (m, 5H), 2.47 – 2.33 (m, 12H), 2.32 – 2.22 (m, 3H), 2.16 (d, *J* = 10.7 Hz, 6H), 2.10 – 1.36 (m, 19H), 1.66 (d, *J* = 5.5 Hz, 3H), 1.33 (d, *J* = 9.5 Hz, 3H), 1.25 (d, *J* = 12.3 Hz, 3H), 1.16 (d, *J* = 6.1 Hz, 3H), 1.08 – 1.04 (m, 4H), 1.01 (s, 3H), 0.93 – 0.84 (m, 2H), 0.27 (s, 1.5H), 0.26 (s, 1.5H).

HPLC chromatogram (λ = 254 and 361nm):

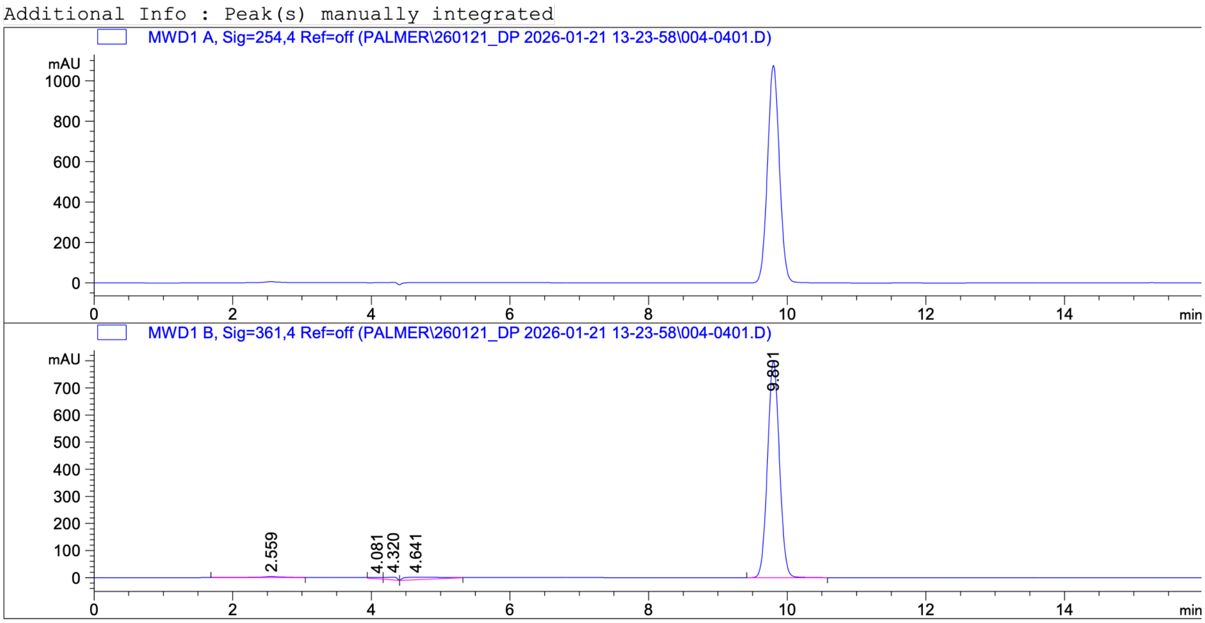

### HRMS spectrum:

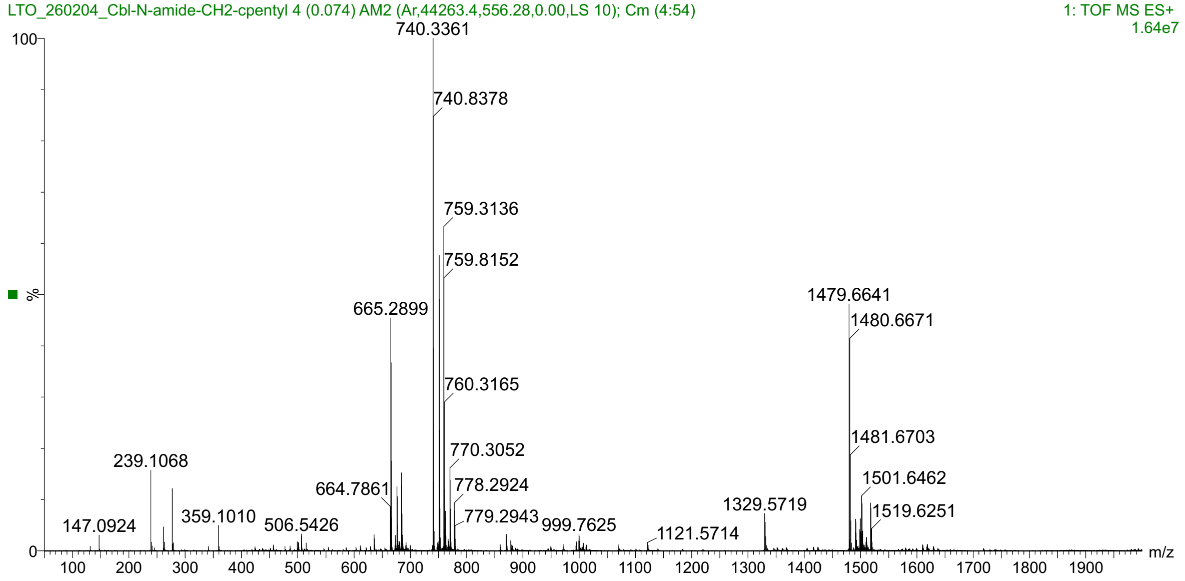


### $^1H$ NMR spectrum (400 MHz, DMSO-$d_6$):

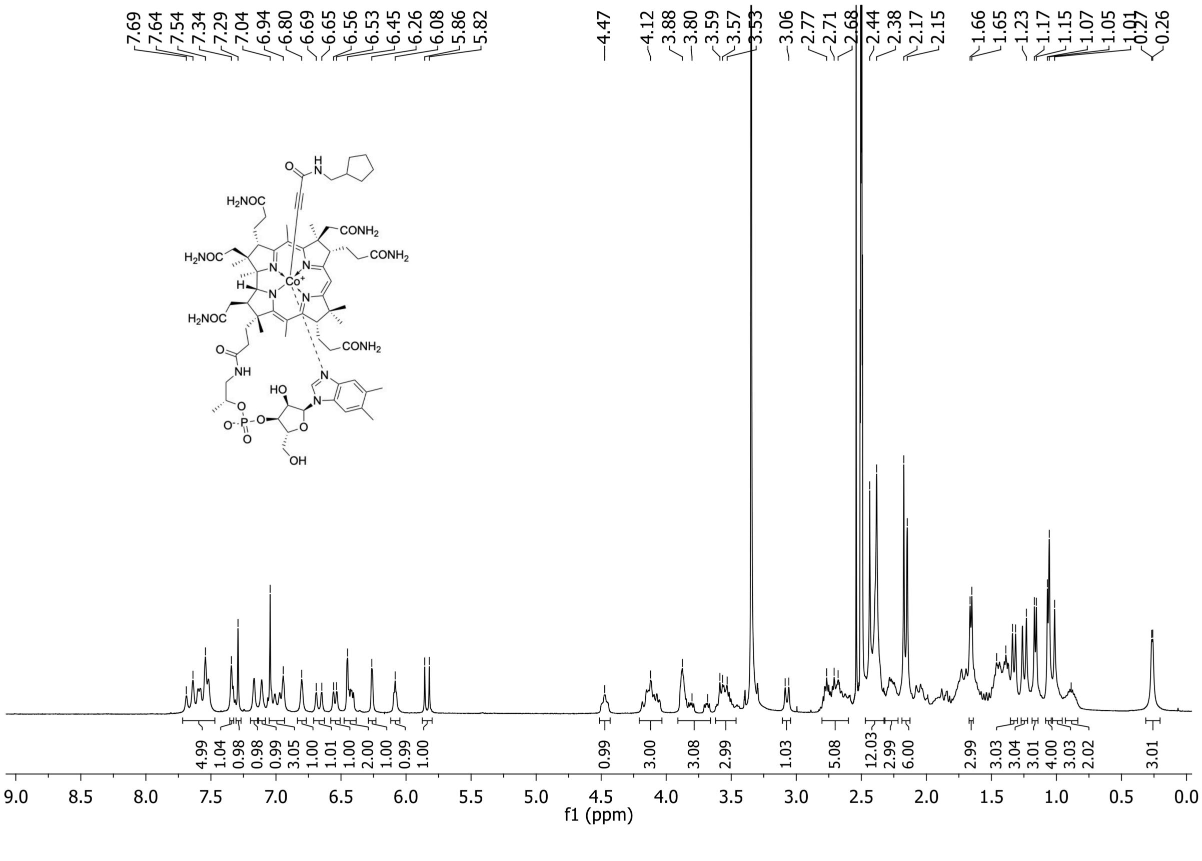

Cbl **71**: Yield: 15.9 mg (48%); HRMS (ESI) m/z $[M + H]^+$ calcd for $C_{71}H_{101}CoN_{14}O_{15}P^+$, 1,479.6635 Da; found, 1,479.6641 Da; $t_R$ (RP-HPLC): 9.69 min; $^1$H NMR (400 MHz, DMSO-$d_6$): δ 7.72 – 7.48 (m, 5H), 7.34 (s, 1H), 7.29 (s, 1H), 7.22 (s, 0.5H), 7.20 (s, 0.5H), 7.17 (s, 1H), 7.11 (s, 1H), 7.05 (s, 0.5H), 7.04 (s, 0.5H), 7.03 – 6.90 (m, 2H), 6.80 (s, 1H), 6.68 (s, 0.5H), 6.66 (s, 0.5H), 6.55 (s, 0.5H), 6.54 (s, 0.5H), 6.45 (s, 1H), 6.44 – 6.38 (m, 1H), 6.26 (d, *J* = 2.3 Hz, 1H), 6.09 (d, *J* = 2.7 Hz, 1H), 5.87 (s, 0.5H), 5.82 (s, 0.5H), 4.51 – 4.44 (m, 1H), 4.24 – 4.06 (m, 3H), 3.90 – 3.68 (m, 3H), 3.61 – 3.49 (m, 3H), 3.08 (d, *J* = 10.3 Hz, 1H), 2.75 – 2.58 (m, 4H), 2.47 – 0.83 (m, 21H), 2.48 – 2.35 (m, 10H), 2.30 – 2.23 (m, 2H), 2.20 – 2.11 (m, 6H), 1.66 (d, *J* = 4.8 Hz, 3H), 1.33 (d, *J* = 16.5 Hz, 3H), 1.24 (s, 3H), 1.17 (s, 3H), 1.10 – 1.01 (m, 6H), 1.01 – 0.87 (m, 4H), 0.27 (s, 1.5H), 0.26 (s, 1.5H).

HPLC chromatogram (λ = 254 and 361nm):

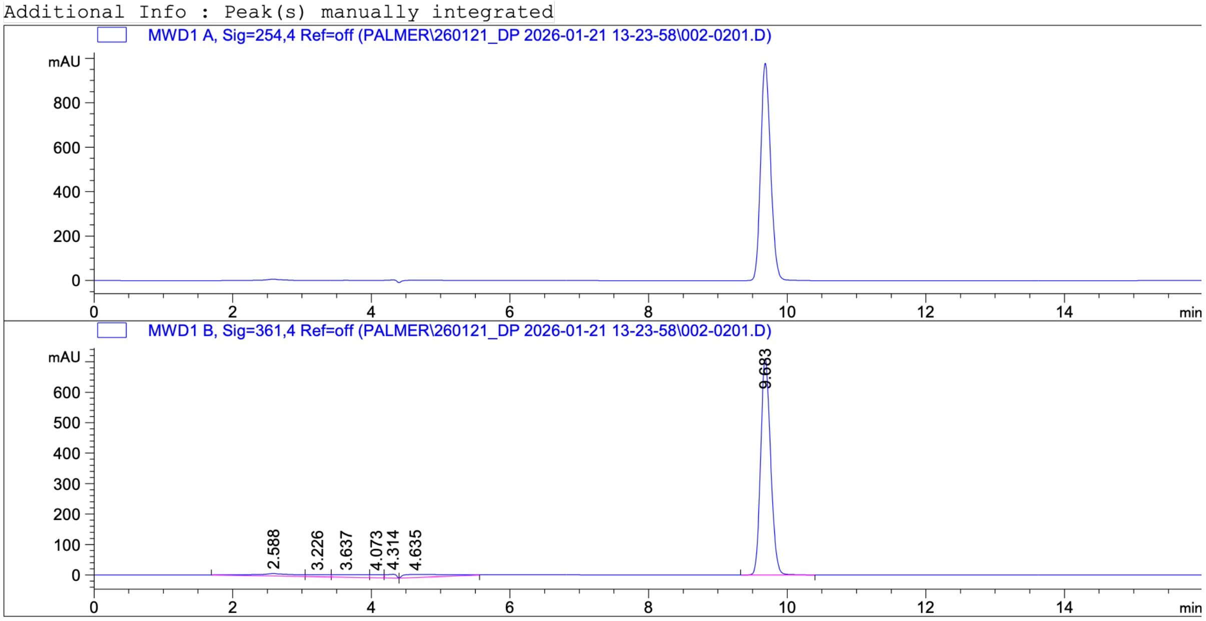

### HRMS spectrum:

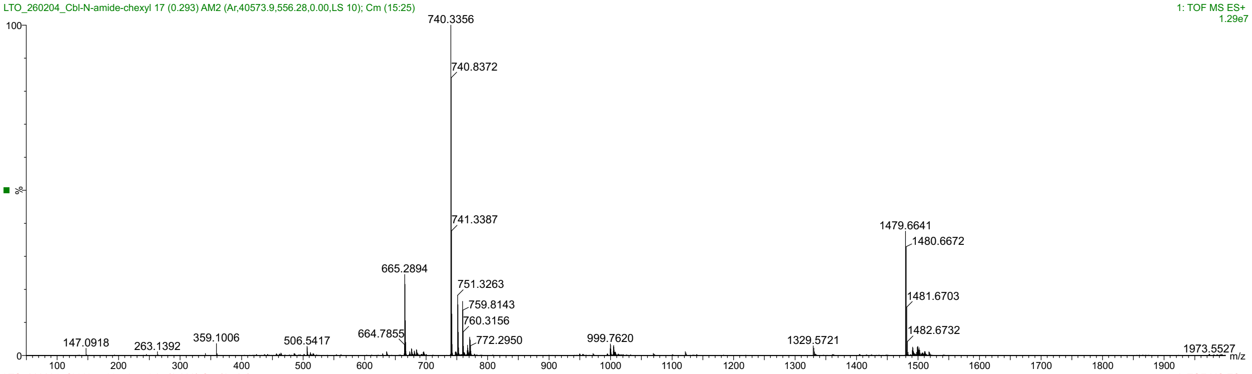


### [1]H NMR spectrum (400 MHz, DMSO-$d_6$):

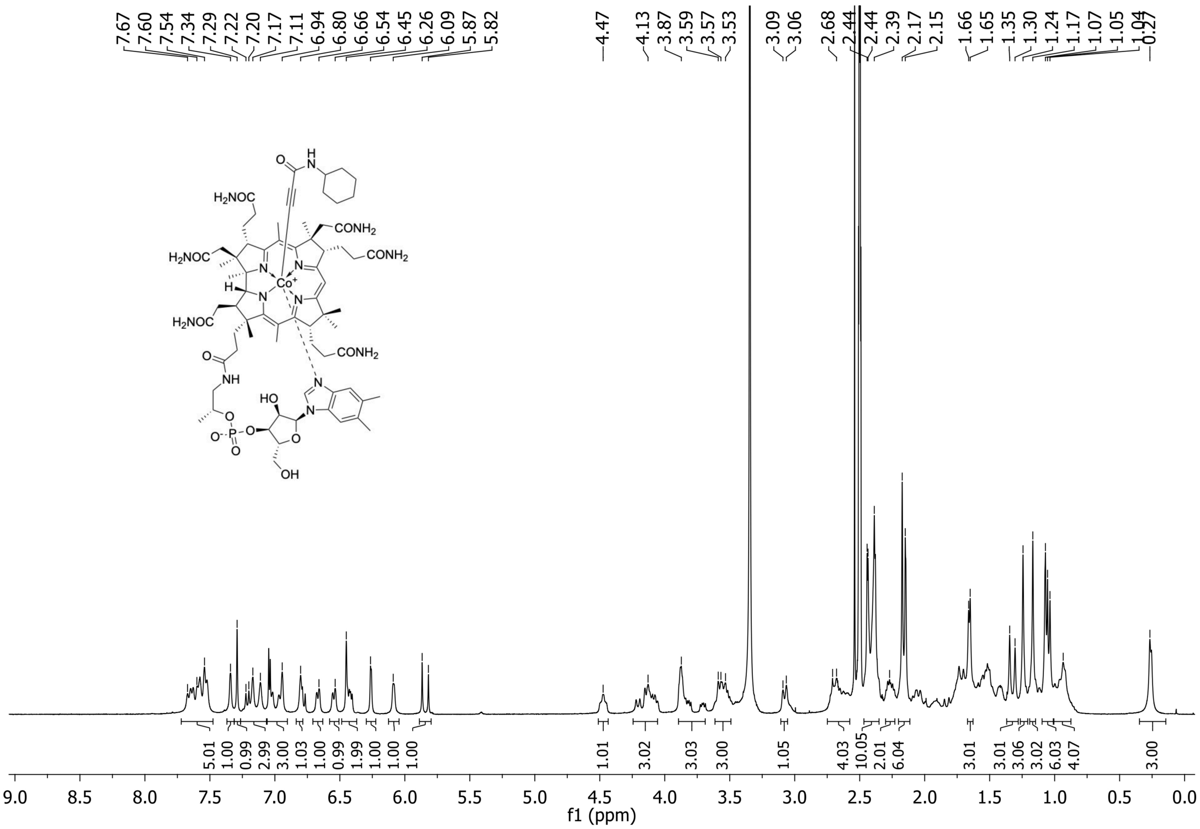

Cbl **72**: Yield: 17.7 mg (54%); HRMS (ESI) m/z [M + H]$^+$ calcd for $C_{72}H_{103}CoN_{14}O_{15}P^+$, 1,493.6797 Da; found, 1,493.6807 Da; $t_R$ (RP-HPLC): 10.11 min; $^1$H NMR (400 MHz, DMSO-$d_6$): δ 7.74 – 7.46 (m, 5H), 7.34 (s, 1H), 7.29 (s, 1H), 7.17 (s, 1H), 7.11 (s, 1H), 7.04 (s, 1H), 7.02 – 6.93 (m, 2H), 6.80 (s, 1H), 6.69 (s, 0.5H), 6.65 (s,0.5H), 6.56 (s, 0.5H), 6.53 (s, 0.5H), 6.453 (s, 0.5H), 6.446 (s, 0.5H), 6.43 – 6.38 (m, 1H), 6.26 (s, 1H), 6.09 (s, 1H), 5.86 (s, 0.5H), 5.82 (s, 0.5H), 4.52 – 4.42 (m, 1H), 4.19 – 4.00 (m, 3H), 3.90 – 3.66 (m, 3H), 3.61 – 3.48 (m, 3H), 3.07 (d, *J* = 10.7 Hz, 1H), 2.75 – 2.60 (m, 5H), 2.45 – 2.35 (m, 10H), 2.32 – 2.21 (m, 3H), 2.20 – 2.12 (m, 6H), 2.11 – 1.38 (m, 19H), 1.66 (d, *J* = 4.9 Hz, 3H), 1.33 (d, *J* = 9.2 Hz, 3H), 1.25 (d, *J* = 13.1 Hz, 3H), 1.16 (d, *J* = 3.8 Hz, 3H), 1.11 – 0.98 (m, 9H), 0.94 – 0.86 (m, 1H), 0.71 – 0.56 (m, 2H), 0.26 (s, 3H).

HPLC chromatogram (λ = 254 and 361nm):

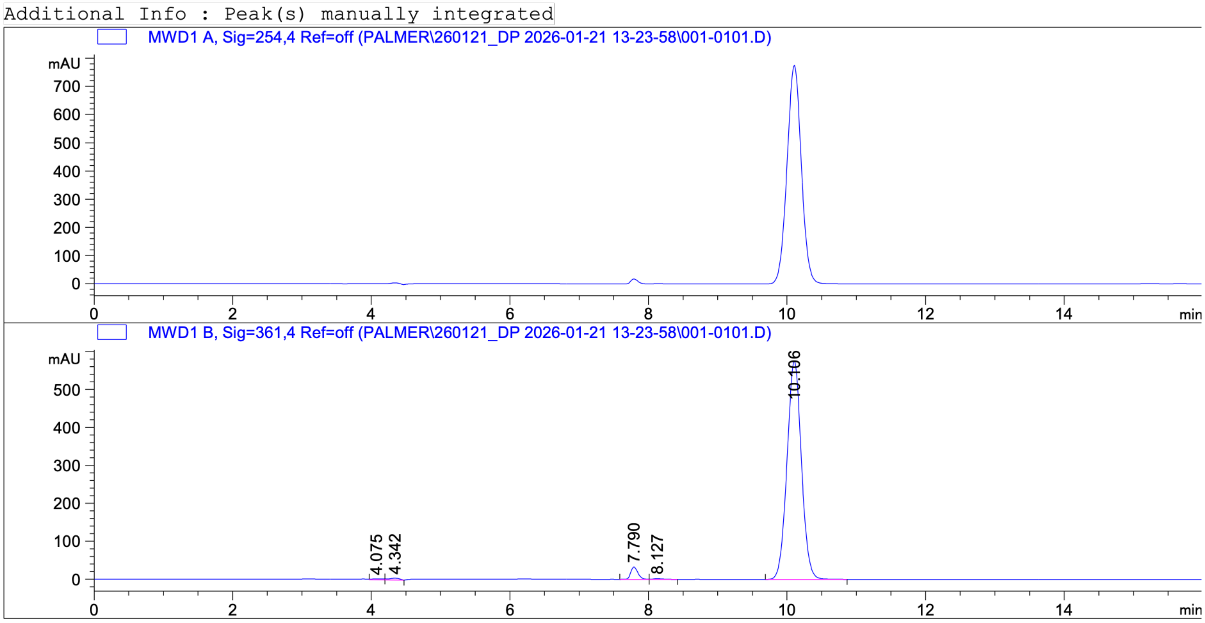

HRMS spectrum:

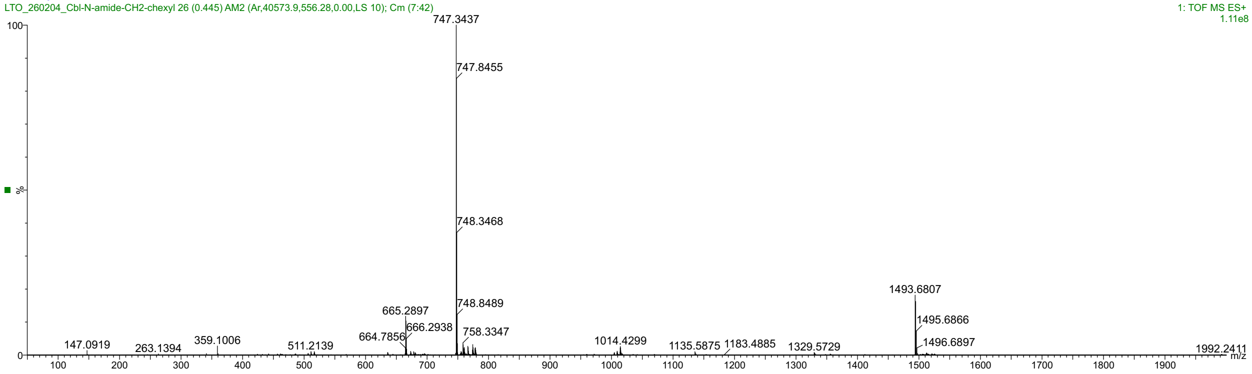


$^1$H NMR spectrum (400 MHz, DMSO-$d_6$):

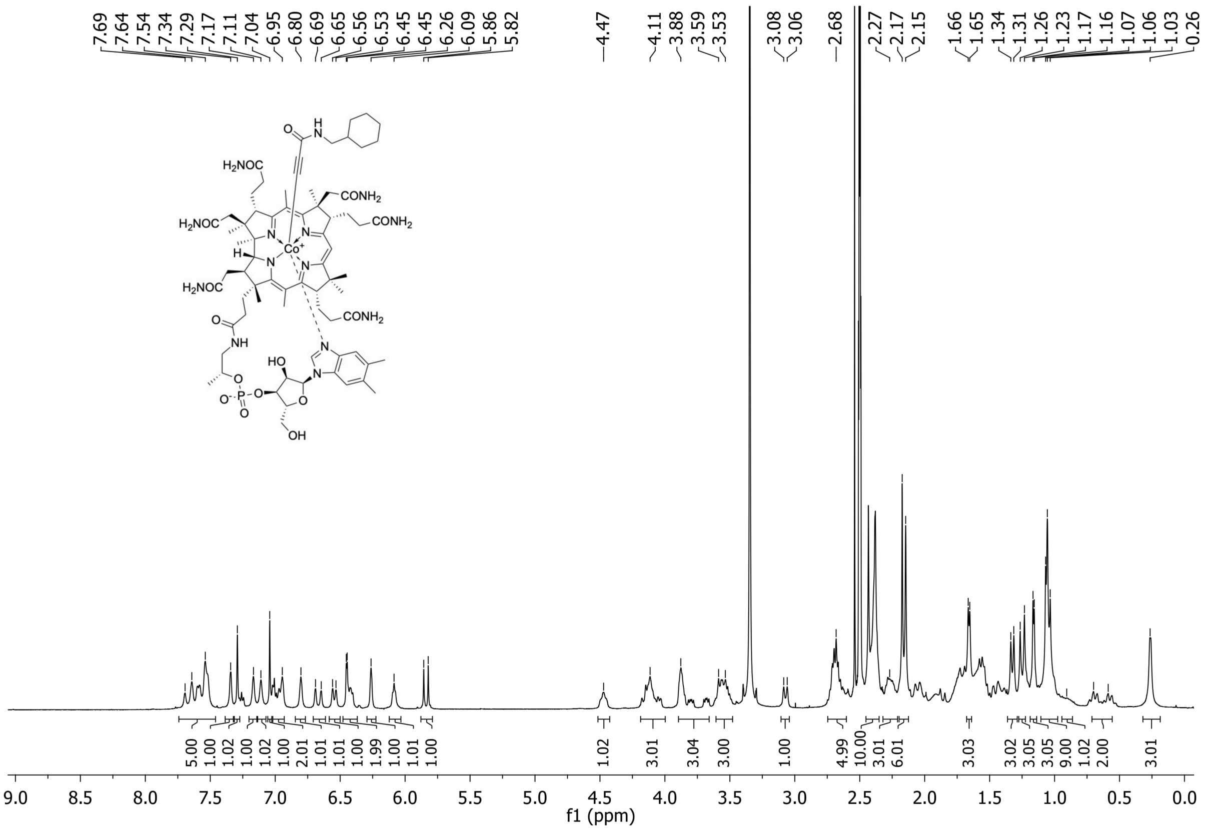

Cbl **73**: Yield: 19.0 mg (58%); HRMS (ESI) m/z $[M + H]^+$ calcd for $C_{72}H_{97}CoN_{14}O_{15}P^+$, 1,487.6322 Da; found, 1,487.6339 Da; $t_R$ (RP-HPLC): 9.58 min; $^1H$ NMR (400 MHz, DMSO-$d_6$): δ 7.78 – 7.51 (m, 6H), 7.35 (s, 1H), 7.29 (s, 1H), 7.26 – 7.11 (m, 5H), 7.09 – 6.92 (m, 5H), 6.80 (s, 1H), 6.69 (s, 0.5H), 6.65 (s, 0.5H), 6.56 (s, 0.5H), 6.53 (s, 0.5H), 6.47 (s, 0.5H), 6.45 (s, 0.5H), 6.43 – 6.37 (m, 1H), 6.26 (s, 1H), 6.11 – 6.03 (m, 1H), 5.83 (s, 0.5H), 5.80 (s, 0.5H), 4.52 – 4.45 (m, 1H), 4.23 – 4.03 (m, 5H), 3.91 – 3.83 (m, 2H), 3.78 – 3.70 (m, 1H), 3.59 – 3.49 (m, 3H), 3.09 – 2.94 (m, 1H), 2.75 – 2.58 (m, 4H), 2.46 – 2.35 (m, 10H), 2.31 – 2.22 (m, 3H), 2.19 – 2.12 (m, 6H), 2.11 – 1.49 (m, 11H), 1.66 (s, 3H), 1.36 – 1.17 (m, 9H), 1.12 – 1.01 (m, 6H), 0.98 – 0.86 (m, 1H), 0.71 (s, 1H), 0.27 (s, 3H).

HPLC chromatogram (λ = 254 and 361nm):

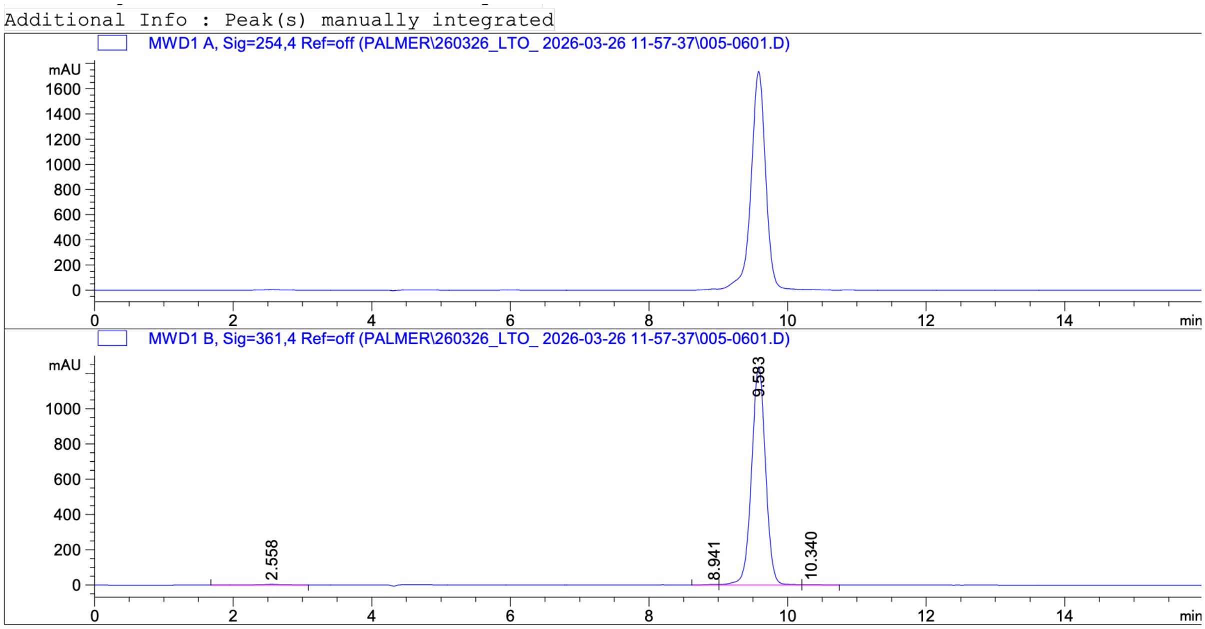

### HRMS spectrum:

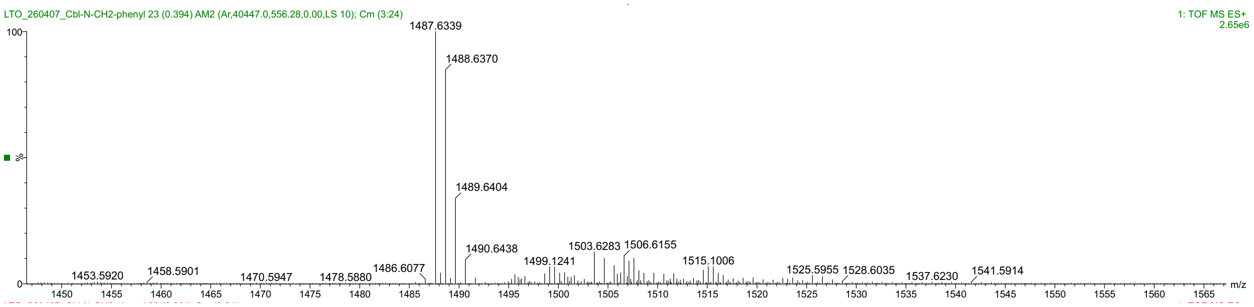


### $^{1}$H NMR spectrum (400 MHz, DMSO-$d_6$):

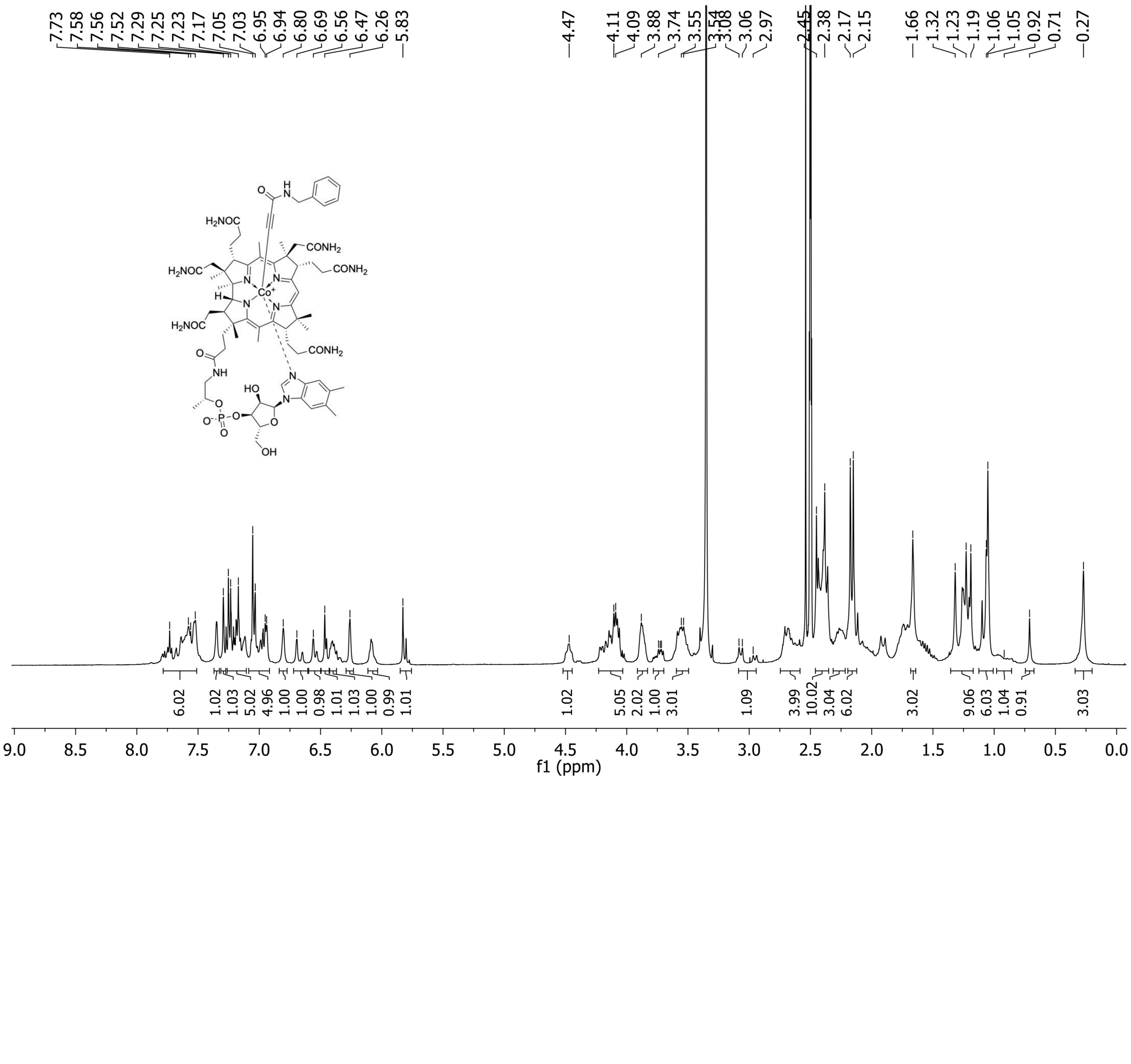
